# A model of working memory for latent representations

**DOI:** 10.1101/2021.02.07.430171

**Authors:** Shekoofeh Hedayati, Ryan O′Donnell, Brad Wyble

## Abstract

Visual knowledge obtained from our lifelong experience of the world plays a critical role in our ability to build short-term memories. We propose a mechanistic explanation of how working memories are built from the latent representations of visual knowledge and can then be reconstructed. The proposed model, Memory for Latent Representations (MLR), features a variational autoencoder with an architecture that corresponds broadly to the human visual system and an activation-based binding pool of neurons that binds items’ attributes to tokenized representations. The simulation results revealed that the shapes of familiar items can be encoded and retrieved efficiently from latents in higher levels of the visual hierarchy. On the other hand, novel patterns that are completely outside the training set can be stored from a single exposure using only latents from early layers of the visual system. Moreover, a given stimulus in working memory can have multiple codes, representing specific visual features such as shape or color, in addition to categorical information. Finally, we validated our model by testing a series of predictions against behavioral results obtained from WM tasks. The model provides a compelling demonstration of how visual knowledge yields compact visual representation for efficient memory encoding.

## Introduction

In the study of memory, working memory (WM) is thought to be responsible for temporarily holding and manipulating information. This capacity to control information is thought to be a keystone of our ability to perform complex cognitive operations. Characterizing WM is an integral part of the birth of cognitive psychology, as decades of research have centered on the question of discovering the capacity and nature of this short-term memory system (e.g., Miller, 1956; when the term *short-term memory* was favored).

One of the central issues in many discussions over the structure of WM is how it is affected by previously learned knowledge (i.e., long-term memory; Baddeley & Hitch, 1974; Cowan, 1988; Cowan, 2019; Ericsson & Kintsch, 1995; Norris, 2017; Oberauer, 2009). Knowledge that emerges from long-term familiarity with particular shapes or statistically common featural combinations enables us to recognize and remember complex objects (i.e., the prototypical shape of a car, or the strokes that comprise a digit). It is widely acknowledged that such information is crucial for building WM representations (Cowan, 1999; Brady, Konkle & Alvarez, 2009; Oberauer, 2009) but there has been little attempt, if any, to mechanistically implement the role of visual knowledge in WM models in spite of abundant behavioral research in this domain (Alvarez & Cavanagh, 2004; Chen & Cowan, 2005; 2009; Hulme, Maughen & Brown, 1991; Ngiam, et al., 2019; Ngiam, Brissenden & Awh, 2019; Yu et al., 1985; Zhang & Simon, 1985; Zimmer & Fischer, 2020). For instance, Alvarez & Cavanagh (2004) demonstrated that the number of items stored in WM is affected by stimulus complexity, with particularly poor performance for Chinese characters. Zimmer & Fischer (2020) expanded on this by showing that the difficulty in remembering Chinese characters is specific to individuals who are not readers of the language. That is to say, the WM capacity for Chinese characters is higher if observers have already been trained on those stimuli. Moreover, Brady, Stormer & Alvarez (2016) have demonstrated that evident memory capacity for natural images is higher than memory capacity for simple colors, as natural images are the stimuli that people have had the most visual experience with compared to artificial shapes. These results can be extended to verbal memory, as performance on immediate recall of a list of words is limited by the number pre-learned chunks represented in long-term knowledge (Chen & Cowan, 2005; Hulme, Maughan & Brown, 1991; Hulme et al., 2003).

Even prior to these findings, there has been extensive theoretical discussion of the necessity to link WM to long-term memory representations. Among the earliest memory schemes was the Atkinson & Shiffrin (1968) model of memory in which representations in long-term memory could be transferred to a short-term storage if needed. Later, Baddeley and Hitch (1974) proposed the multicomponent model of WM. In this model the short-term storage for visual information (i.e., visuospatial sketchpad) was shown to be dependent on visual semantics and episodic long-term memory with a bidirectional arrow indicating the flow of information (Baddeley, 2000). This idea is also carried by theories of activated long-term memory account (Cowan 1988, 1999, 2001; Ericsson & Kintsch, 1995) which is also discussed by Oberauer (2009). In this account (also known as the embedded process framework), WM representations are built by activating pre-existing representations within the long-term memory.

The above accounts (i.e., multiple components and activated long-term memory) provide a venue toward a WM mechanism integrated with long-term knowledge, but their lack of computational specificity has made it challenging to make testable predictions of *how* knowledge reflects in WM mechanisms. This includes addressing questions such as “How do we form rapid memories of novel configurations (Lake et al., 2011)?” and “Why is WM capacity higher for familiar items (Yu et al., 1985; Zhang & Simon, 1985; Zimmer & Fischer, 2020)?”

To fill this gap, we implemented a computational WM model in conjunction with a visual knowledge system named Memory for Latent Representations (MLR). The proposed model simulates how latent representations of items embedded in the visual knowledge hierarchy are encoded into WM depending on their level of familiarity. Subsequently, the encoded items in WM can be retrieved by reactivating those same latent representations in the visual knowledge system. This paper outlines a candidate model for storing and retrieving visual memories of complex shapes in a dedicated pool of neurons and provides empirical validation of the flexibility of WM.

### The new MLR model of working memory

The MLR model takes advantage of recent innovations in generative deep learning models to represent and reconstruct visual stimuli that are embedded in the visual knowledge. Rather than storing unidimensional attributes as in other recent working memory models (Bouchacourt & Buschman, 2019; Schneegans & Bays, 2017; Lin & Oberauer, 2019; Swan & Wyble, 2014), MLR can encode and reconstruct arbitrary shape attributes, such as a particular handwritten digit, or an article of clothing. To achieve this, MLR uses the latent distributions from the hidden layers in a pre-trained deep neural network. In this context, *latent* is a representation of a stimulus attribute such as the shape, color, or category of a stimulus. Figure 1 illustrates an example of digits that can be reconstructed from a simple two-dimensional latent space in a variational autoencoder (i.e., VAE) trained on the MNIST dataset, which is a collection of 60,000 hand-written digits (Kingma & Welling 2013).

**Figure 1.**
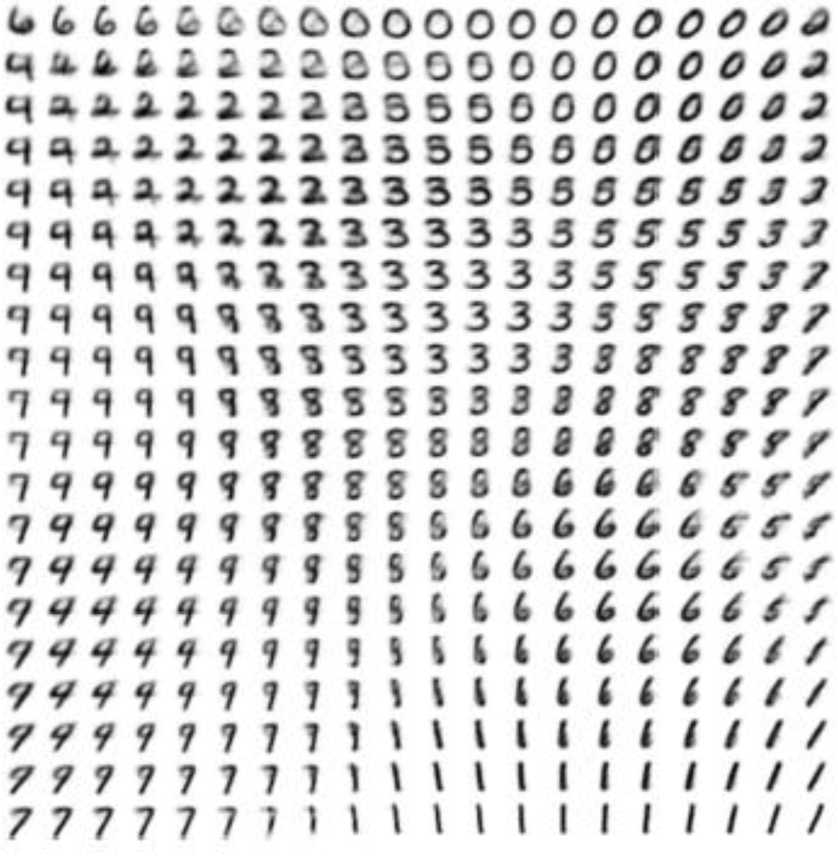
A latent space for MNIST digits from Kingma & Welling (2013). These digits represent the output of a generative model when a particular x,y location in the latent space is activated and then used to drive a reconstruction back to a full 28×28 pixel image of a digit.

### The visual knowledge in MLR

In the context of our work, knowledge is the emergent feature of a trained sensory system. For instance, the connectivity of the visual system is adjusted via experiencing visual objects in a child, which results a visual knowledge system. MLR captures two fundamental characteristics of visual knowledge that enables us to explain its interaction with WM. In the proposed framework, these two features are compression of visual information and categorical representations. Later, we show how these aspects of visual knowledge would affect WM capacity and precision of retrieved items for familiar and novel items (Brady, et al., 2008; Yu et al., 1985; Zhang & Simon, 1985; Zimmer & Fischer, 2020).

#### Compression

The amount of visual sensory input that we receive at every moment is enormous. Therefore, efficient data compression is essential given the limited-resources available to the perceptual system. It is likely that the visual ventral system represents familiar visual patterns with fewer neurons at successively later levels of the pathway (i.e., LGN, V1, V2, V4, IT) of the visual system (Bates & Jacob, 2020). Hence, hierarchical visual knowledge can be formed from the compression of visual data (Norris & Kalm, 2020; Ngiam, Brissenden & Awh, 2019) by *learning,* via synaptic plasticity (Lamprecht & LeDoux, 2004), to encode and decode that data with high visual specificities. In this framework, later levels of the ventral stream (i.e., IT cortex) can represent specific shape patterns with minimal loss of visual details relative to earlier layers (i.e., V1) despite utilizing a smaller number of neurons to form the representation. This is due to connections between neurons encoding feature conjunctions in a hardwired fashion (VanRullen, 2009). Importantly, this compression is only effective for representations that are deeply familiar to the visual system (i.e., in which there have been thousands of exposures, sufficient to develop perceptual expertise, see Pelli, Burns, Farell & Moore-Page, 2006) and not for novel stimuli.

Consistently, empirical data has shown long-term memory to have highly detailed representations of visual objects (Brady, et al., 2008; Konkle et al., 2010; also see Experiment 2).

#### Categorical representation

A familiar object also can have a conceptual/categorical representation. It has been demonstrated that whenever we perceive patterns that correspond to familiar concepts, that information is rapidly interpreted by the visual system to be translated into categorical codes that exist at an abstract level (Huang & Awh, 2018; Potter & Faulconer, 1975, Potter, Valian & Faulconer,1977; Potter, 2018). For instance, for an experienced reader of the Roman alphabet, take note of the character: ‘A’. This character can be represented as a series of strokes with fine grained visual details that include the rightward slant, or the conceptual representation of A in a form of purely categorical information. The memory of seeing the familiar letter ‘A’ could be composed of one or both of these codes depending on task requirements.

The framework described here as visual knowledge, entailing the compression and categorical representation for visual data endows WM with a doubly efficient representation of familiar objects. In other words, familiar objects benefit from compressed representations of visual information and also abstract categorical codes as they are encoded into WM, whereas novel objects lack such efficient representations. Figure 2 illustrates the diagram of hypothetical compressed and categorical representations of a handwritten digit ‘5’ as it is being processed by the visual system. The key point here is that with increasing depth into the ventral stream the visual character is represented by progressively fewer neurons but the loss of detail is minimal as the stimulus category is familiar to the visual system. Moreover, this visual representation elicits a separate categorical representation which is even more compact than the visual representation, though it lacks the visual data.

**Figure 2.**
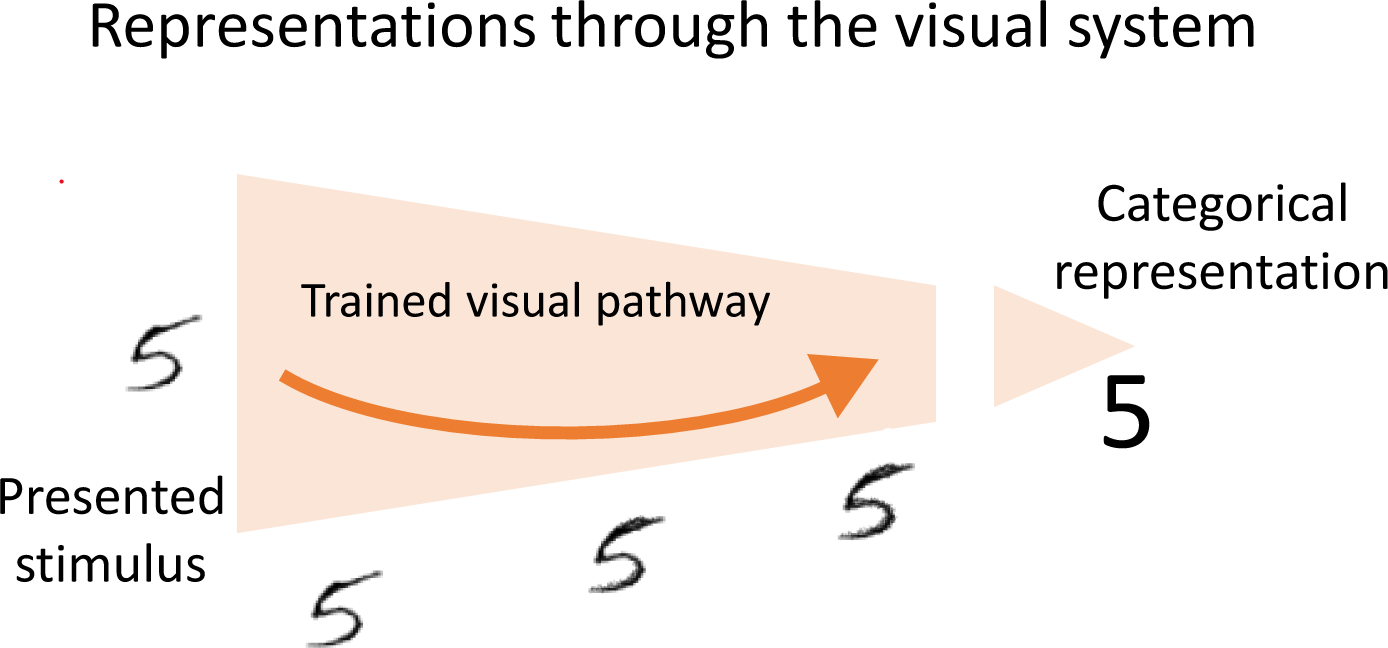
The compression and categorical representation of a single stimulus. The trained visual pathway represents the stimulus with specific visual details in all layers with little loss of visual specificities. The width of the cone reflects the number of neurons involved in the representation at different stages of processing. The final representation at the highest level would elicit a categorical representation that lacks the visual information.

### Informal Description of MLR

Memory formation in MLR can *rapidly* and *selectively* encode specific attributes (e.g., shape, color, label) from one or more visual items within a distributed neural representation using a tokenized binding pool (Swan & Wyble, 2014). In MLR, information is encoded with varying levels of efficiency depending on the degree to which it matches representations embedded within the pre-trained visual knowledge hierarchy. The visual knowledge is built using gradient descent to train a modified VAE on a set of stimuli combining handwritten digits and articles of clothing (MNIST, LeCun, 1998; and fashion-MNIST, Xiao, Rasul & Vollgraf, 2017). We chose to build our model based on a fully connected VAE rather than a more complex convolutional network, because it is simple in terms of layers, and generates smooth latent spaces. More complex models would provide more detailed reconstructions, but our goal is to develop a clear and explainable theory rather than an optimized memory system.

In a trained VAE, familiar stimuli are encoded efficiently into a small-dimensional latent space (analogous to IT cortex) and classified into categorical labels, while novel stimuli can only be represented into higher-dimensional latents closer to the beginning of the visual pathway (analogous to V1 cortex).

As illustrated in Figure 3, a visual stimulus is processed by the feedforward portion of the visual knowledge system to produce compressed shape and color representations of the object as well as a categorical label of each attribute. A binding pool stores a representation of selected features and/or labels according to a set of tunable parameters. These parameters control the proportion of different kinds of information that flow from the knowledge hierarchy into the memory trace.

**Figure 3.**
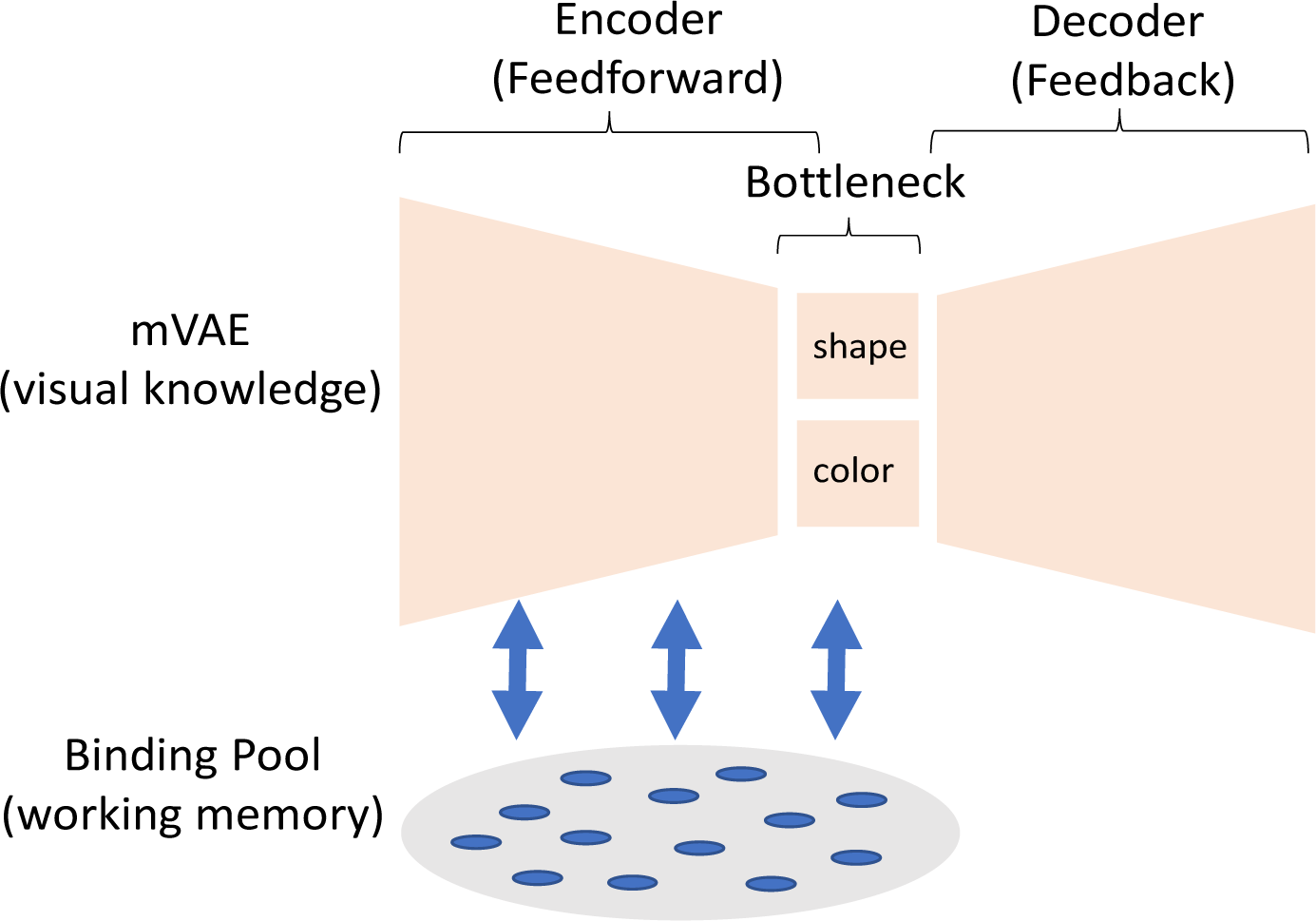
The simplified architecture of MLR with its two major elements: visual knowledge as represented by the modified VAE and the working memory as represented by the binding pool. We modified the bottleneck to represent shape and color in separate maps.

Each memory trace binds visual forms, colors, and categorical labels together into a single tokenized representation that can be stored alongside other tokenized representations of objects.

### A brief description of the landscape of WM theories

A clarifying assumption of MLR is that the memories are stored in a particular group of neurons that are allocated specifically to the role of memory storage and sits apart from the sensory areas themselves (Figure 4a). This account is in accordance with classic theories of prefrontal cortical involvement in WM (Goldman-Rakic,1995; Miller, Erickson & Desimone,1996). This can be contrasted with models that imply distinct representations for visual and non-visual forms of memory (Baddeley & Hitch, 1974; Figure 4b), and embedded process models that distribute the storage of information through a variety of memory and sensory systems (Cowan 1988, 1999; Cowan, Morey, & Naveh-Benjamin, 2020; Morey, 2018; Pasternak & Greenlee, 2005; Teng & Kravitz, 2019; Figure 4c).

**Figure 4.**
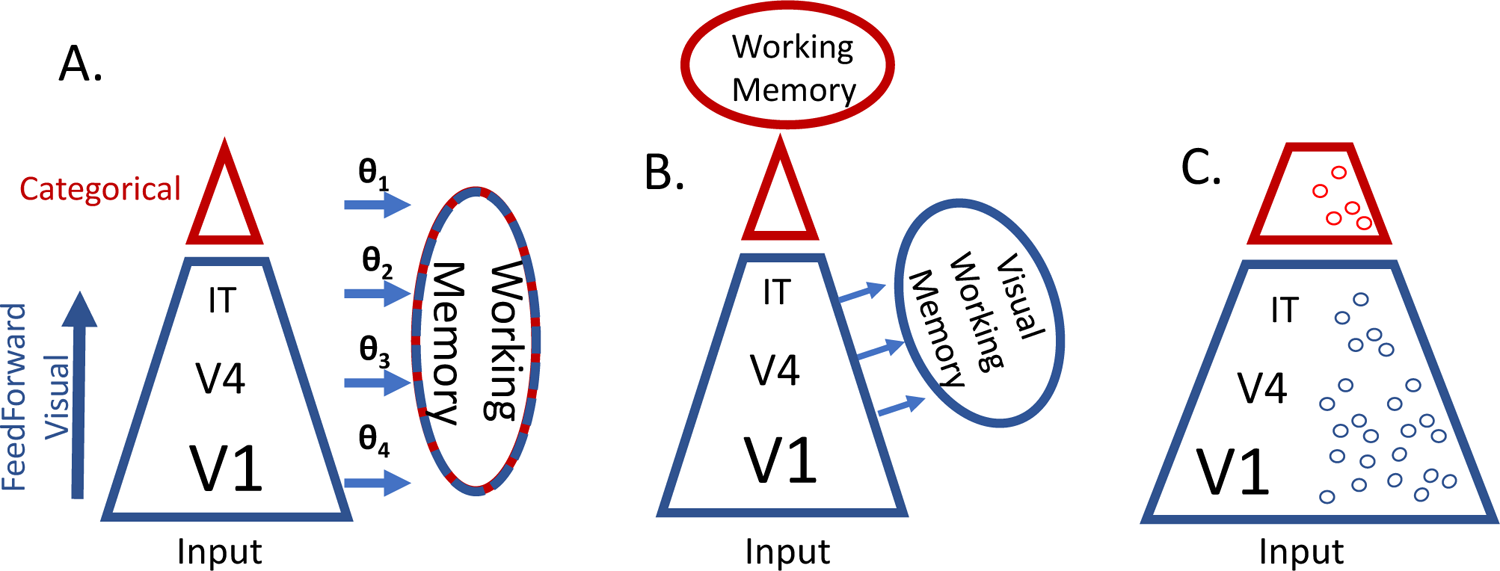
Three architectures for working memory as it relates to the visual system. **A.** The MLR model as proposed here has a single memory representation that encompasses visual and categorical information in varying proportions according to task-dependent tunable parameters. **B.** A working memory model that suggests there are distinct systems for maintaining visual and non-visual forms of information **C.** A working memory model that has its representations embedded in the sensory system

Much thought and experimental evidence has been allocated to adjudicating between these hypothetical architectures (Cowan, 1999, 2001; Lee, Kravitz & Baker, 2013; Logie, Camos & Cowan, 2020; Morey, 2018). Our goal is not to refute competing accounts at this point, but rather to provide a possible functional implementation of how knowledge *could* be combined with WM. We are advancing one particular instantiation of such an account, and will discuss a comparison between approaches in the discussion. However, it should be emphasized that the mechanism of storing latent representations in a binding pool as described here has some generality and could be adapted to other architectures (i.e., it would be easy to use two binding pools, one for visual and one for non-visual information). A primary goal of this paper is to clarify potential implementations to develop computational formalisms for comparison of different architectures.

### Modelling Philosophy of this account

Computational models can be used in a variety of ways to advance psychological theories (Guest & Martin, 2021). In this case, the MLR model provides a new conceptualization of WM via implementation that achieves a range of benchmarks, some of which are functional and others of which are neural. This can be considered as an abductive approach, in which a likely explanation is proposed for a set of data. We consider the problem of WM models to exist in the M-open class (Clarke, Clarke & Yu, 2013), which means that though it is impossible to specify the exact biological system, we are able to distill useful constraints from behavior and biology.

### Functional constraints of MLR

To motivate our account, we start with a list of empirical constraints that define the relevant functional capabilities of memory and several key neural plausibility constraints.

MLR is constrained by empirical data obtained from human behavior, termed *functional constraints.* To minimize the complexity of the model’s ancillary assumptions, these constraints will be met in a qualitative fashion rather than by matching of specific empirical data points.

*Reconstructive:* Although reconstruction of visual stimuli is not required in typical visual memory tasks, it is a form of retrieval, and people are able to draw or otherwise reconstruct the specific shape of remembered objects, particularly if subtle visual details need to be remembered (Bainbridge, Hall & Baker, 2019; Carmichael, 1929; Kosslyn, 1995). The MLR accounts for this form of retrieval through pixel-wise reconstruction of images.

*Multiple codes*: A memory of a familiar stimulus can be represented by a variety of different codes, from low level visual details to abstract categorical information (Potter & Faulconer, 1975; Potter, Valian & Faulconer, 1977; Potter, 2017). The MLR can store a combination of different kinds of information from a single stimulus, including its visual details or its categorical labels.

*Encoding Flexibility*: Within the visual memory of a single item, specific attributes (e.g., color, shape, etc.) are stored with varying degrees of precision. That is, depending on the task, some visual features are encoded more accurately than others. For example, Swan, Collins and Wyble (2016) showed that memories for features that are relevant to the task (e.g., color of an oriented bar) are remembered more precisely than irrelevant features (e.g., orientation of the bar).

Parameters in MLR control the ratio of distinct attributes of an object that are stored in memory. In the simulations here, color and shape are treated as distinct attributes but in a larger model, the set of tunable attributes could include any stimulus dimension for which there are distinct representations. This would be a separate latent space in the context of an VAE, or a separate cortical area in the context of neuroscience (Konkle & Caramazza; 2013)

*Representing Novel stimuli*: WM performance is more efficient for previously learned items (Yu et al., 1985; Zimmer & Fischer, 2020), however, humans can still encode novel configurations that they have not seen before (Lake et al., 2011; see also Experiment 1). Similarly, MLR stores and retrieves novel shapes that it has not seen before, although those memory reconstructions are less precise compared to shape categories that the model was trained on.

*More Efficient representations of familiar items:* Frequently-experienced objects drawn from long-term knowledge have compressed representations with high visual detail (Brady et al., 2008; Konkle et al., 2010). This allows more familiar objects to be stored in memory compared to novel stimuli (Hue & Erickson, 1988; Zimmer & Fischer, 2020). MLR achieves this by encoding compressed representation of familiar items generated by a smaller number of neurons, whereas it resorts to encoding features represented in larger number of neurons if the object is unfamiliar.

*Individuated Memory for Multiple items:* Memory for one visual display or trial is able to store several different items and retrieve them individually using a variety of cues. For example, one could store a series of colored shapes and then retrieve a specific item based on one particular attribute, such as color, shape, location or serial order. In addition, people can store repetitions of the same item as well as the temporal order of different items (see Swan & Wyble, 2014 for more detailed discussion of this point, see also Bowman & Wyble, 2007; Kanwisher, 1991; Mozer, 1989 and Swan & Wyble, 2014). MLR uses tokens as pointers to individuate different items, including repetitions.

*Content Addressability and Binding:* Memory representations include a form of binding in which multiple attributes can be attached to one another, which is often thought of as object bindings when those attributes belong to distinct visual objects. Also, *content addressability* means that such bindings can be used to retrieve any attribute associated with the object from any other attribute. For example, a colored, oriented line can be accessed either through its color or orientation (Gorgoraptis et al., 2011). MLR can use any attribute or code associated with a token to retrieve other information from that token.

*Mutual interference:* Storing multiple items and/or additional features of an item (i.e., shape, color, size, etc.) in WM causes interference that degrades the memory precision based on the number of items stored (Wilken & Ma, 2004) and the number of attributes within one stimulus (Swan, Collins & Wyble 2016). The shared neural resources in MLR cause overlapping representations to interfere with one another, both for attributes within a stimulus and between stimuli.

### Neural constraints of MLR

*Rapid encoding and forgetting:* WM representations are thought to be the result of persistent neural activity (Compte, et al., 2000; Fuster & Alexander, 1971) or transient latent synaptic representations that can be rapidly changed (Rose et al. 2016; Szatmáry & Izhikevich, 2010). These representations allow for rapid encoding and removing of visual information. The shared binding pool of MLR stores visual information by creating temporary activity states in a matrix that is intended as a generalization of either a population of self-sustaining neural attractors or other forms of rapidly modifiable representations (e.g., silent synapses; Rose et al. 2016). These activations are received via fixed randomly assigned weights from different layers of the visual knowledge hierarchy.

*Hierarchical structure of ventral stream:* The ventral visual pathway contains at least part of the visual knowledge that is gradually formed through extensive experience with the world. In primates, this pathway has cells that vary along a spectrum from receptive fields that are tuned to orientation and color in the earliest layers such as LGN and V1 cortex, up to cells that have large receptive fields and that are tuned to more complex shapes such as faces and complex configurations (Grill-Spector, Kourtzi & Kanwisher; 2001; Kanwisher, McDermott & Chun, 1997). In MLR, visual knowledge is based on a VAE architecture as illustrated in Figure 5. The VAE resembles the hierarchical structure of the visual ventral stream with more generic representations at the early level and more compressed representations at higher levels (i.e., the bottleneck) with the number of neurons decreasing progressively. In a VAE, the layers from the bottleneck to the output translates between latent representations and a pixelwise representation. These are similar to the extensive feedback projections that extend backwards down the ventral stream from higher to lower order areas (Bullier, 2001; Lamme, Super & Spekreijse;1998).

**Figure 5.**
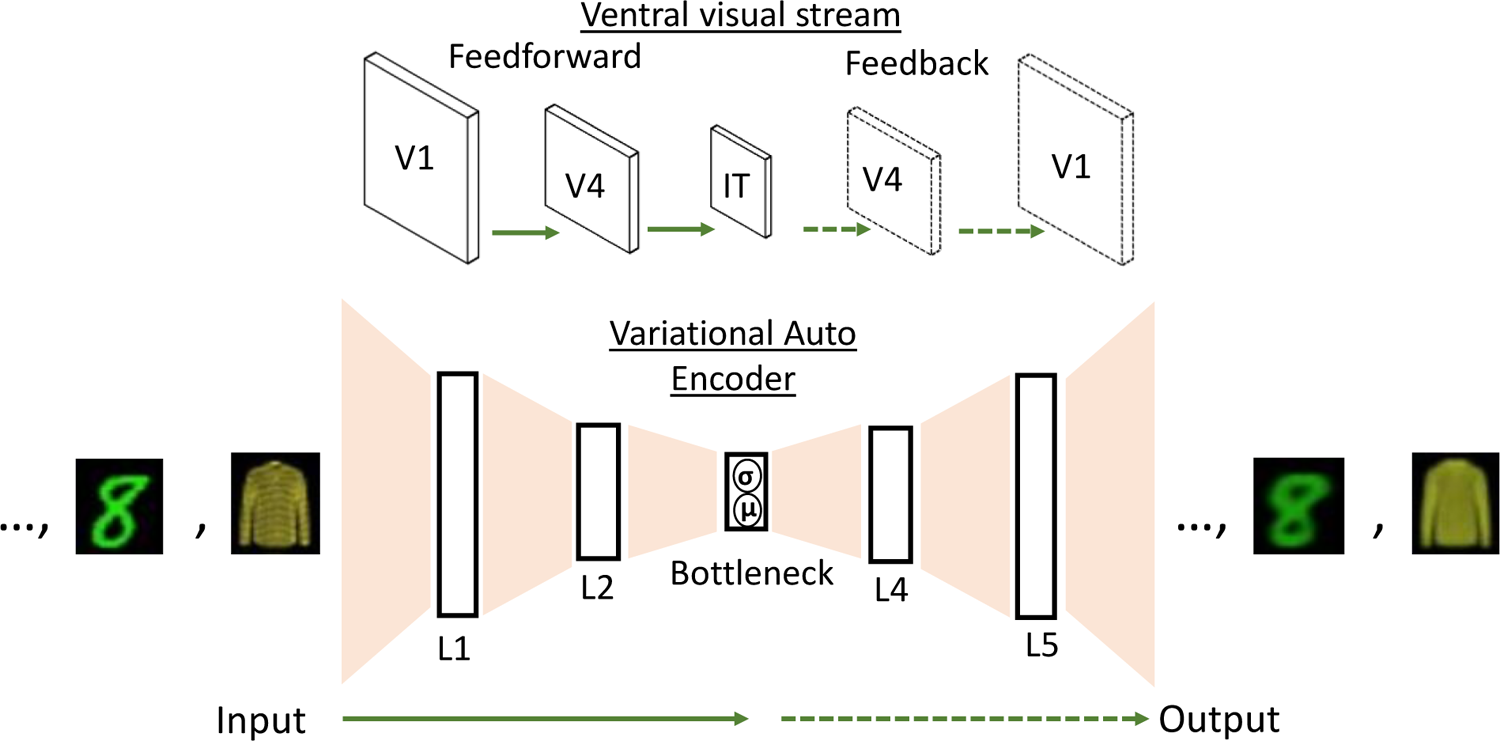
Illustration of the architecture of a VAE (Kingma & Welling 2013) and its coarse neuroanatomical correspondence. In the neuroanatomical projection, solid arrows correspond to feedforward connections from V1 to IT cortex (or L1 to bottleneck in the VAE) and dashed arrows refer to feedback projections in the reverse direction (or from bottleneck to L5 in the VAE). The inputs were either colorized versions of MNIST and f-MNIST. Note that one image at a time is fed into the VAE.

*Training through synaptic weight adjustments:* Our biological brain is an ever-changing system that alters its connectivity over time to represent the statistical regularities of the environment. Likewise, MLR learns statistical regularities underlying visual categories through experiencing abundant exemplars that are used during the training phase. As a result of this training, the network’s connectivity is tuned to better reconstruct the visual stimuli from compressed representations in the bottleneck. Moreover, training in the VAE occurs without explicit labels or supervision, akin to how a child can learn to see through exposure to patterned information.

### The Architecture of MLR

The model is composed of two components: a modified variational autoencoder (mVAE) operating as visual knowledge and a binding pool (BP), the memory storage that holds one or more objects (see Figure 6 for the detailed architecture).

**Figure 6.**
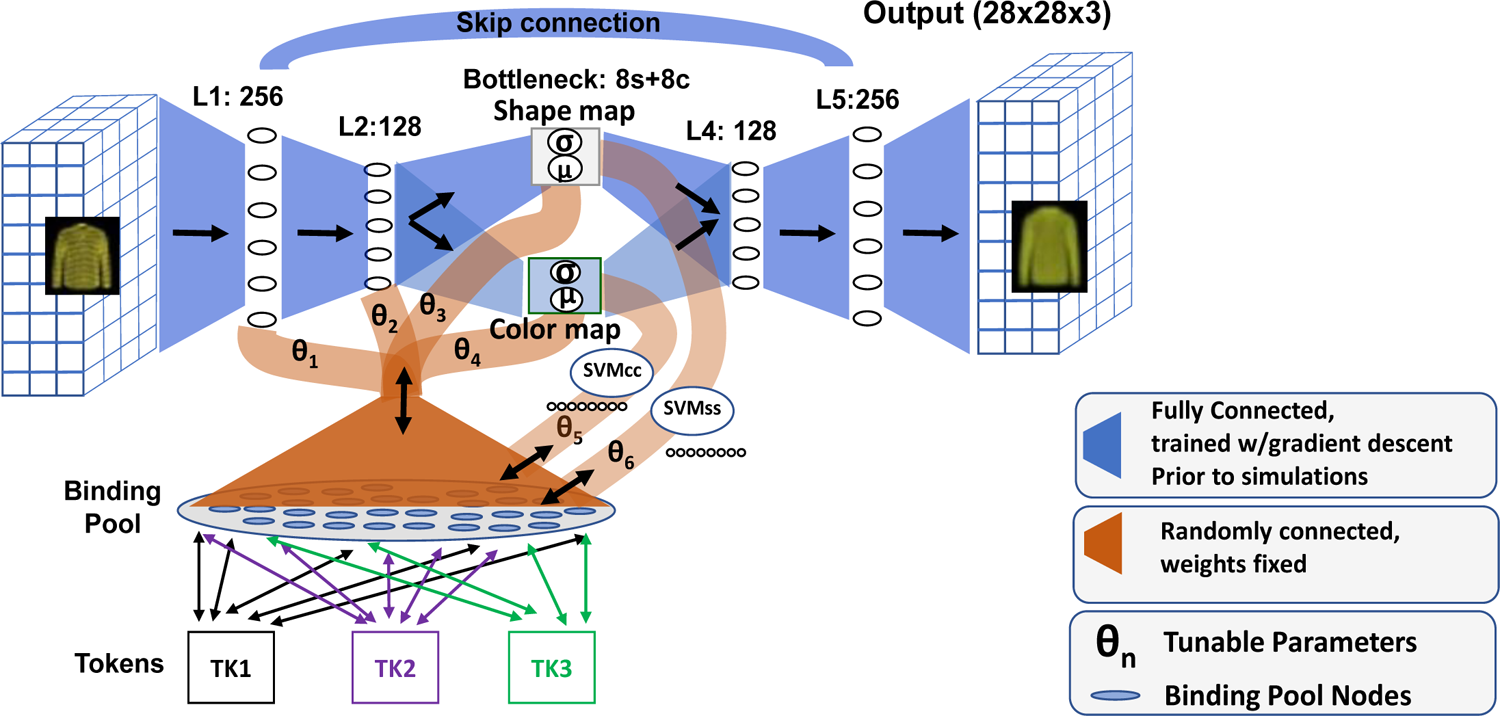
The complete MLR architecture that consists of the mVAE, binding pool, tokens, and classifiers for extracting labels (SVMss and SVMcc). Information flows in only one direction through the mVAE but can flow bidirectionally between the latent representations and the binding pool. Tokens are used to differentiate individual items. Note that three tokens are shown here but there is no limit to the number of tokens that can be allocated.

*mVAE*: The VAE (Kingma & Welling 2013) is an hourglass shape fully connected neural network consisting of three main elements – feedforward, bottleneck and feedback – which are trained by using a colorized variant of MNIST (LeCun, 1998) and fashion-MNIST (Xiao, Rasul & Vollgraf; 2017) stimulus sets prior to any memory storage simulations. We modified the VAE by dividing the bottleneck into two separate maps – a color map and a shape map – to represent each feature distinctively. See the appendix for details about the colorized stimuli and the modified objective functions.

*Feedforward pathway*: Translates information from a pixel representation into compressed latent spaces as a series of transitions through lower dimensional representations. This is typically called the *encoder* in autoencoder models.

*Shape and Color maps:* Typically, the bottleneck layer of a VAE that has the smallest number of neurons consists of one map. To generate distinct feature maps, we divided the bottleneck into two separate maps: one for representing shape and the other one for representing color. Each of the two maps is fully connected to the last layer of the feedforward pathway and the first layer of the feedback pathway.

*Feedback pathway*: Translates information from the compressed shape and color maps into pixel representations as a series of transitions through progressively higher dimensional representations. This is typically called the *decoder* in autoencoder models.

*Skip Connection:* To allow reconstruction of novel stimuli without involving the shape and color maps, a skip connection was added to the mVAE that linked the first layer to the last layer.

Anatomically, this would be the equivalent of a projection between layers with V1 cortex (Thomson 2010)

*Categorical labels:* In order to apply categorical labels to a given stimulus, we used a standard support vector machine classifier (SVM; Cortes & Vapnik, 1995). The SVM maps representations in the latent spaces onto discrete labels for different stimulus attributes such as shape or color. See appendix for the details of the SVMs.

*Binding Pool (BP):* The BP uses a modified formulation of the model described in Swan & Wyble (2014) and is similar to a Holographic Reduced Representation (Plate, 1995). It is a one-dimensional matrix of neurons that is bidirectionally connected to each layer of the feedforward pathway (L_1_, L_2_, shape and color maps) as well as the outputs of the SVM classifiers which provide one-hot categorical labels of shape and color. The BP stores a combined representation of the information from each of these sources for one or more stimuli in individuated representations indexed by tokens. The bidirectional connections allow information to be encoded into the BP, stored as a pattern of neural activity, and then projected back to the specific layers of the mVAE to produce selective reconstruction of the encoded items. The connection between the BP and the latents is accomplished through normally generated, fixed weights.

These are not trained through gradient descent but are assigned at the beginning of the simulation for a given model.

*Tokens:* The tokens function as object files (Marr 1976; Kahneman & Triesman, 1984) for each specific stimulus (e.g., token 1 stores stimulus 1). Having tokens allows multiple items to be stored, even if they are spatially overlapping. They index representations that are held in the BP and do not store information about the individual stimuli. See the appendix for additional details on the tokens.

### The MLR implementation

*Architecture:* The mVAE consists of 7 layers. Input layer (L_i_; dim= 28 x 28 x 3), Layer 1 (L_1_; dim= 256), Layer 2 (L_2_; dim= 128), bottleneck (color map, dim= 8; shape map, dim = 8), Layer 4 (L_4_; dim= 256), Layer 5 (L_5_; dim= 256) and the output layer (L_o_; dim=28 x 28 x 3). A skip connection was added from L_1_ to L_5_. The BP layer is connected to the feedforward layers of mVAE bidirectionally (Figure 6, also see appendix).

*Datase*t: Training was done using the MNIST stimulus set (LeCun et al., 1998) consisting of 70,000 images of 10 categories of digits (0-9) and fashion-MNIST set (i.e., f-MNIST; Xiao, Rasul & Vollgraf, 2017) which has the same number of images as MNIST but for 10 categories of clothing (T-shirt/top, Trouser, Pullover, Dress, Coat, Sandal, Shirt, Sneakers, Bag and Ankle boot). To add additional attributes to the dataset, we colorized all images using 10 distinct colors – red, blue, green, purple, yellow, cyan, orange, brown, pink, teal – with minor variations (see Figure 7 for examples and appendix for details of the color values).

**Figure 7.**
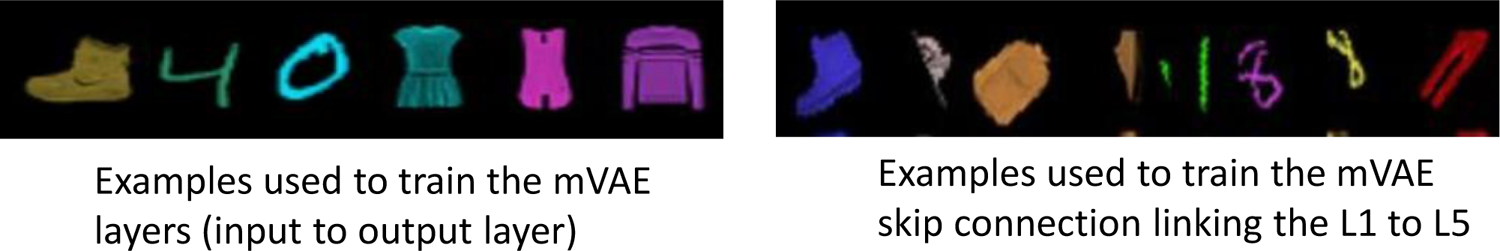
Colorization of MNIST and Fasion-MNIST inputs using 10 prototypical colors with independent random variations on the RGB channels. Left: images used to train the mVAE. Right: Transformed images of the same dataset that were used to train the skip connection.

*Training and testing the mVAE:* The mVAE was trained on 120,000 images with 200 epochs (batch size= 100) with three objective functions to the train shape, color and the skip connections distinctively. See appendix for details on objective functions and training.

*BP memory encoding of latents*: Once the mVAE was trained, memories could be constructed by shifting information from the latent spaces into the BP with 2500 neurons in total. The effective number of neurons representing each item was 1000 since 40% of the BP was allocated to each token. Such memories are constructed with a matrix multiplication of the activation values of a given latent space (i.e., L_1_, L_2_, shape and color map) or one-hot categorical labels, by a randomly generated and fixed (i.e., untrained by gradient descent), normally distributed set of weights. This multiplication produces a vector of activation levels for each neuron in the BP. Multiple attributes can be combined into one representation in the BP by summing the activation values from multiple encoding features and then normalizing them. Equation 1 demonstrates the encoding of activations in the BP, where B_β_ represents each node in the BP, N_t,β_ represents the connection matrix between the BP nodes and the token, X_f_ represents the activations in a given latent space, n is the number of neurons in the latent space that is being stored in the BP, and L_f,β_ is the connection matrix between the latent space and the BP.

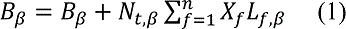

*BP memory encoding of categorical labels:* The color and shape category labels estimated by an SVM classifier, as an analog of categorical representations, could also be encoded into the BP. The shape labels were extracted from SVM_SS_ (i.e., an SVM trained to decode shape labels from the shape map) and the color labels were extracted from SVM_CC_. (i.e., an SVM trained to decode color labels from color map). Shape was one-hot coded in a vector of length 20 (10 digits and 10 fashion items), while color used a vector of length 10. Either or both of these vectors could be added to a BP representation through matrix multiplication described above. Reconstructions from the BP were converted into a one-hot vector with a max function.

*One-shot encoding of novel shapes in BP:* Novel shapes were 6 examples of colorized Bengali characters. The colorization of Bengali characters was similar to that of MNIST and f-MNIST. The colored novel images were used as inputs to the model, and activations from L_1_ and shape and color maps were encoded and retrieved from the BP to compare the efficiency of encoding from these layers (Figure 10).

*Storing multiple items:* Working memory, though limited in capacity, is capable of storing multiple objects. The tokens in MLR provide an index for each object that can be later retrieved from the BP as in Swan & Wyble (2014). Without such an indexing mechanism, more than one object cannot be stored in the memory. Each token contacts a random, fixed proportion of the binding pool, effectively enabling those units^1^ for memory encoding while that token is active. Any number of tokens can be stored in this way, although more interference is expected as the number of stored tokens increases. This mechanism enables multiple distinct sets of attributes to be stored in each token, effectively binding those attributes into one object. The tokens can be retrieved individually and in any order. Once stored in this way, a token can reactivate its portion of the BP to reconstruct the attributes associated with it. Moreover, tokens enable content addressable recall in that a given attribute (e.g., the shape or color of a digit) can be used as a retrieval cue to determine which of several tokens was associated with that specific attribute.

Then, that token can be activated to retrieve the other attributes associated with it (see Swan & Wyble, 2014 for more details).

*Memory Maintenance in BP:* The binding pool is a simple implementation of a persistent-trace model that holds the vector of activation produced by the encoding operation(s). This is consistent with self-excitatory neural attractors, or silent synaptic storage (Rose et al. 2016). The specific mechanism of trace-maintenance was not a crucial question in this implementation as there was no time course or delay of activity over time.

*Token Retrieval:* To determine which token was linked to a given visual form (e.g., a shape map representation), information can be passed from a given latent through the BP to determine which token has the strongest representation of that particular latent. Equation 2 illustrates the retrieval of a given token Z_t_.Other parameters are similar to that of Equation 1.

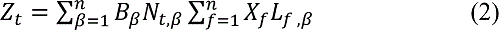

*Memory Reconstruction and model’s evaluation:* Memory reconstructions to any given latent or one-hot vector were accomplished by retrieving the associated token and multiplying the entire BP vector by the transpose of the same fixed weight matrix that was used during the encoding of that representation. As represented by Equation 3, the result is a noisy reconstruction of the original latent activity state, which can be processed by the rest of the mVAE in the same manner as visual inputs.

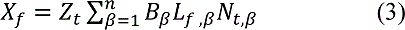

Two methods were used to evaluate the quality of memory reconstructions of MLR. 1) Representations in the shape and color maps were classified by radial basis SVMs, which were trained to decode shape (one of 20) or color (one of 10) using the remaining 10,000 MNIST and 10,000 fashion MNIST as test set stimuli. The classification allowed us to assess the amount of shape and color information in the shape and color maps before and after memory reconstruction. Note that we also used the same pre-trained classifiers to create the labels and to assess memory performance 2.) An alternative measure of the accuracy of reconstructing the original image was to correlate the reconstructed pixels with the original stimulus. We used this approach to quantify reconstructions of novel stimuli which have no pre-learned categories. A detailed implementation of the model was written in Python 3.7 using pytorch toolbox and can be found in https://osf.io/tpzqk/ (The original VAE code that we modified was retrieved from a GitHub repository: https://github.com/lyeoni/pytorch-mnist-VAE/blob/master/pytorch-mnist-VAE.ipynb).

### Simulation results

1. *The mVAE disentanglement prior to memory encoding*: Figure 8 shows reconstructions from shape, color and both maps respectively. The results of classification accuracies in Table 1 show that color and shape representations were successfully disentangled in their corresponding maps. In other words, the shape map contained mostly shape information and the color map represented mostly color. This is a coarse approximation of the general finding that the ventral visual stream has specialization of cortical maps for different types of information (Cohen et al., 2014; Konkle & Caramazza, 2013). The benefit of such anatomical disentanglement in the context of a memory model like MLR is that it permits top-down modulation to select particular kinds of information for promotion to WM because the control signals only need to operate on the scale of selection regions of cortex, rather than individual neurons. That said, the *complete disentanglement* of color and shape as we achieve here is not likely to be a real phenomenon but is very helpful for demonstrating the principles of encoding attributes selectively.

**Figure 8.**
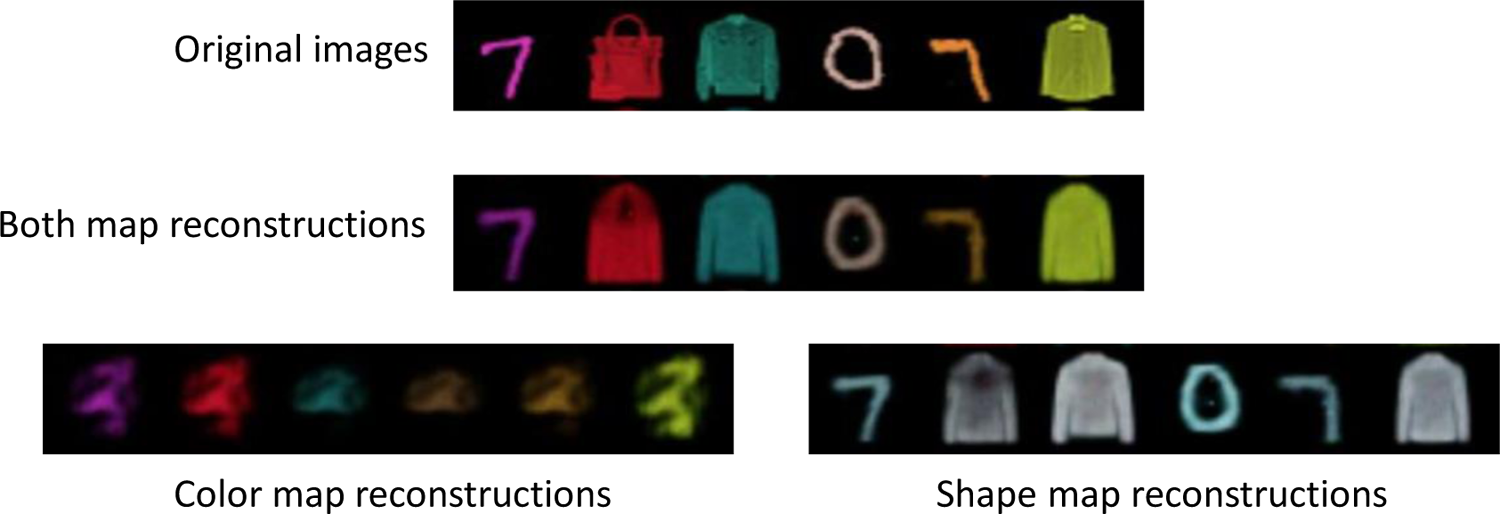
Reconstructions from the mVAE. Information from just one map is shown by setting the activations of the other map to 0. Both maps together produce a combined representation of shape and color, showing that the model is able to merge the two forms of information that are disentangled across the two maps. The model only processes one item at a time in these simulations, and these are combined into single figures for ease of visualization.

**Table 1.** Classifiers accuracies (%) of mVAE for information represented in shape and color maps

|  | Classifier type |  |  |  |
| --- | --- | --- | --- | --- |
|  | SVM<sub>SS</sub> | SVM<sub>SC</sub> | SVM<sub>CC</sub> | SVM<sub>CS</sub> |
| Shape/color maps | 84 (.20) | 22 (.30) | 87 (.40) | 15 (.1) |

Note. The table shows mean classification accuracies (%) for the mVAE for 10 trained models. The values in parentheses represent standard errors. SVM<sub>SS</sub> represents an SVM trained on shape labels using data from the shape map. On the other hand, SVM<sub>SC</sub> was trained to decode color labels from the shape map. SVM<sub>CC</sub> Represents an SVM trained on color labels using data from the color map, whereas SVM<sub>CS</sub> was trained to decode shape labels from the color map. Chance performance is 10% for classifiers trained on color labels (SVM<sub>CC</sub> and SVM<sub>SC</sub>) and 5% for classifiers trained on shape labels (SVM<sub>SS</sub> and SVM<sub>CS</sub>). As shown, SVM<sub>SC</sub> and SVM<sub>CS</sub> accuracies are above chance, but much smaller than SVM<sub>SS</sub> and SVM<sub>CC</sub> respectively.

1. 2. *The BP encoding and retrieval of visual features:* Projecting information from the latent representations into the BP and then back to the mVAE allows us to reconstruct the original activity pattern of that layer. Figure 9 illustrates examples of single items encoded individually and then reconstructed using the mVAE. Table 2 indicates the classifiers’ accuracies averaged across 10 models for determining the shape and color of items according to which layer of the mVAE was encoded and then retrieved. According to the simulation results, it is evident that memory retrieval from shape and color maps is more precise than reconstructions from L_2_ and L_1_. Hence, compression resulted in more accurate memory retrieval, presumably due to the noisier reconstruction of the larger L1 and L2 latents. It is important to note that retrieval process of the familiar shapes requires images to pass through all the subsequent layers including the shape and color maps (e.g., L_2_ representations are stored in the BP, then projected back to L_2_ and then reconstructed by passing through the maps, L_4_, L_5_ and the output layer. Likewise, classifying the accuracy of the memory formed from the L_2_ layer involves reconstructing the L_2_ latent from the BP, then passing it forward to the shape and color maps and classifying those map activations with the SVMs).

**Figure 9.**
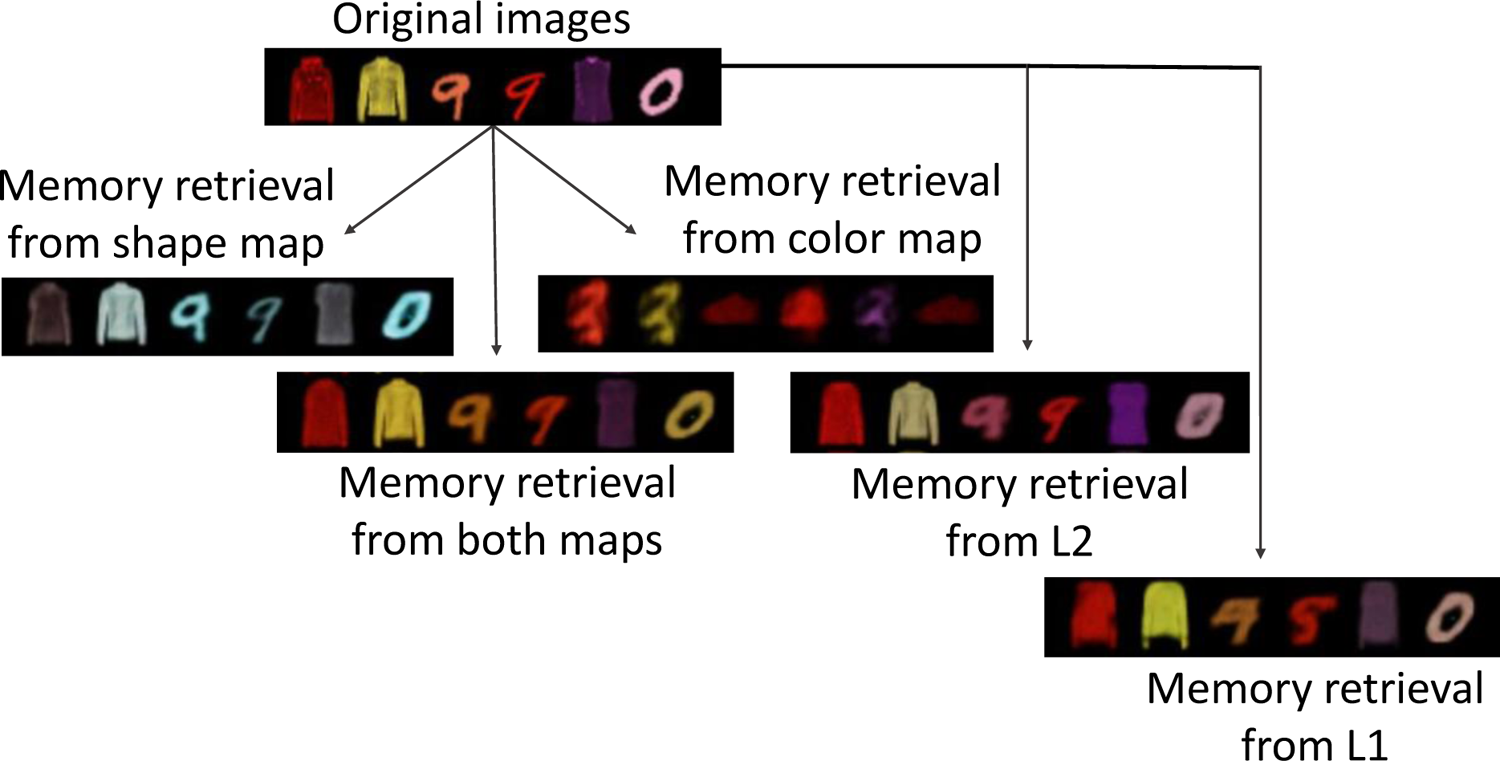
Demonstration of different latents that can be stored from one of the trained models (both shape and color were encoded in all conditions). Note that the reconstructions are visually less precise for memories formed from L_1_ and L_2_ latent spaces compared to the shape and color maps. Each item is stored individually in a separate BP, but the examples are combined into single images for ease of visualization.

**Table 2.** Mean classification accuracy (%) of shape and color information based on encoding conditions

| Encoding conditions | Classifier type |  |  |  |
| --- | --- | --- | --- | --- |
|  | SVM<sub>SS</sub> | SVM<sub>SC</sub> | SVM<sub>CC</sub> | SVM<sub>CS</sub> |
| Shape map and Color map | 83 (.43) | 20 (.36) | 80 (1.45) | 15 (.29) |
| Shape map only | 83 (.36) | 20 (.45) | 9 (.18) | 4 (.22) |
| Color map Only | 4 (.24) | 8 (.69) | 83 (.83) | 15 (.33) |
| L2 | 75 (.55) | 18 (.45) | 69 (2.87) | 14 (.34) |
| L1 | 44 (1.82) | 13 (.85) | 52 (2.96) | 7 (.34) |

Note. The table indicate means of classifier accuracies (%) after memory retrievals of a single stimulus from each layer for 10 independently trained models. The values in parentheses indicate standard errors. Rows correspond to different encoding conditions, showing which latent(s) were stored in the binding pool. Shape map only and color map only indicates that only shape or color of a stimulus was encoded and retrieved. L<sub>1</sub> and L<sub>2</sub> representations were passed forward to the shape and color maps after being stored in the BP to be classified.

1. 3. *Storing multiple attributes and codes of one stimulus:* MLR is able to flexibly store specific attributes of a given stimulus such that BP representations are more efficiently allocated for a particular task. For example, when just color is expected to be task relevant, the BP representation can largely exclude shape information which comes with a small increase in accuracy for reconstructing color. This matches human performance which shows that even when remembering the color of an oriented arrow in WM, there is a small but measurable cost for storing both color and orientation (Swan Collins & Wyble 2016). This is evident in Table 2 wherein the classification accuracy of retrieving color was improved when the shape map was not stored by setting its encoding parameter to zero. The reverse phenomenon was not observed (i.e., shape was not improved by eliminating color; SVM_SS_ in Table 2). It should be noted that the random assignment of BP nodes to each feature map would always result in overlapping activation patterns for different attributes and therefore interference, however the extent of interference depends on the number of BP nodes as well as the number of attributes that are being encoded.

Moreover, MLR is able to store categorical labels of information alongside the visual information in a combined memory trace. By converting the output of a classifier into a one-hot representation, a neural code of label can be stored into the BP, summing with the representations of the shape and color maps. This allows a memory to contain unified categorical and nuanced shape/color information within a single trace (e.g., remembering that one saw a ‘5’ and it had this particular shape or color). As we will show, the categorical labels have shown to be more resistant to interference as more items are being encoded into memory (see Table 3).

**Table 3.** Accuracy of retrieving visual and categorical information (%) as a function of set size

|  | conditions |  |  |  |  |  |  |  |
| --- | --- | --- | --- | --- | --- | --- | --- | --- |
|  | 1s | 1c | 2s | 2c | 3s | 3c | 4s | 4c |
| Set size |  |  |  |  |  |  |  |  |
| 1 | 83 (.56) | 81 (.59) | 82 (.66) | 80 (.82) | 75 (.58) | 82 (.46) | 83 (.46) | 87 (.69) |
| 2 | 63 (.56) | 56 (.79) | 63 (.40) | 56 (.71) | 64 (.53) | 72 (.69) | 81 (.41) | 85 (.41) |
| 3 | 50 (.60) | 45 (.55) | 50 (.69) | 45 (.50) | 53 (.74) | 62 (.36) | 75 (.59) | 80 (.46) |
| 4 | 40 (.65) | 38 (.56) | 39 (.79) | 38 (.65) | 43 (.69) | 54 (.55) | 67 (.49) | 73 (.37) |

Note. Mean classifier accuracy (%) of retrieved items as a function of set size in conditions 1 and 2, and the mean accuracy of one-hot labels before and after storage in memory as a function of set size in conditions 3 and 4. The values in parentheses indicate standard errors computed over 10 independently trained models. In all cases the accuracy declines as more items are stored in memory, however, labels are more resistant to interference as shown in condition 4, especially when the amount of visual information stored in memory decreases.

1. 4. *Encoding of Novel stimuli:* MLR has the ability to store and retrieve novel shapes (i.e., Bengali characters) that it has not seen before. To do so, it stores the L_1_ latent into the BP and retrieves it via the skip connection. The skip connection is critical to reconstruct novel forms, since the nature of the compressed representations in the maps force the reconstructions to resemble familiar shapes (Figure 10).

1. 5. *Encoding multiple visual items:* Tokens allow individuation of different items in memory (Kanwisher, 1991; Mozer, 1989), which in this case occurs by allocating each item to a subset of the binding pool as introduced in earlier work (Bowman & Wyble 2007). Tokens thus provide object-based clustering of attributes in memory. Each item stored in the pool causes interference for the others as their representations partially overlap. Thus, representational quality degrades as additional items are added to memory in agreement with human behavior (Wilken & Ma, 2004).

Note that here we are reconstructing the actual shape and specific colors of the items, not just their categorical designations. As the number of items stored in memory gets larger, the quality of those representations visibly decreases as demonstrated in Figure 11. Note also that the color of the retrieved items becomes more similar as set size increases, reflecting the overlap in representation between the different items. This is emblematic of the interference observed in storing multiple visual stimuli in Huang & Sekuler (2010). To quantify this decrease of memory quality, the number of items stored was increased incrementally and reconstruction accuracy was assessed by classifying the retrieved item (Table 3 and see simulation 6 below).

**Figure 10.**
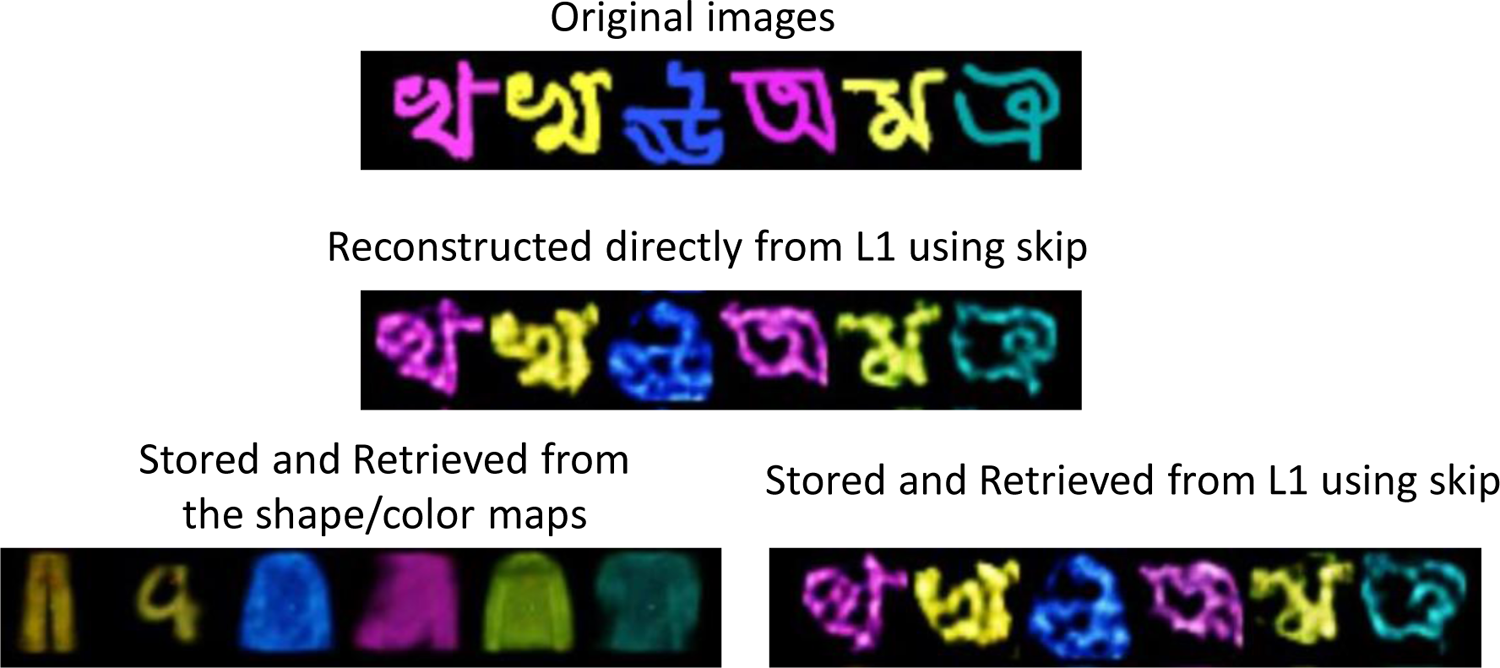
Illustration of storing a single novel Bengali character six times. The original images were reconstructed as familiar shapes when the BP stored the shape and color maps (bottom left). However, the successful reconstructions can be seen when the BP stores the L_1_ activations and retrieve them via the skip connection. These representations are connected in one image for simplicity but each Bengali character was stored and retrieved from an empty memory store.

**Figure 11.**
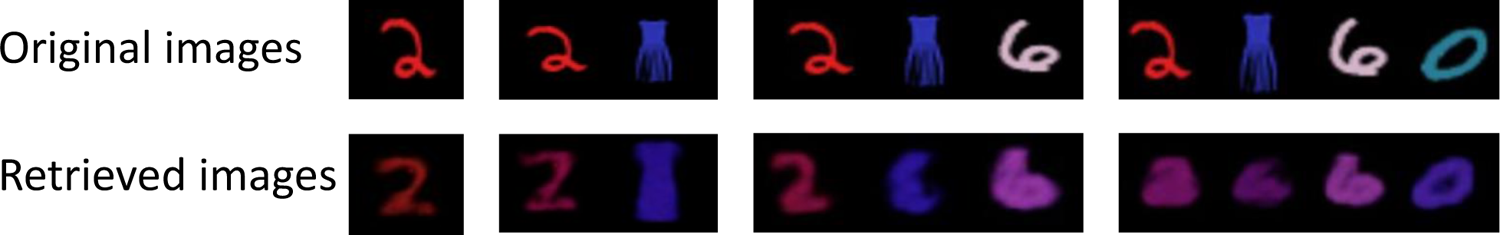
Illustration of the storage and retrieval of 1, 2, 3 and 4 items in memory. The interference increases as more items are stored in the BP. This results in inaccurate reconstructions of both shape and color, as well as the emergence of ensemble encoding.

1. 6. *Multiple codes for multiple objects*: To assess the memory retrievals for visual and categorical information as a function of set size we consider four encoding conditions replicated for set sizes 1-4:

Condition 1 (encode visual, retrieve visual): shape and color map activations are stored together in the BP for each item; Then, either shape is retrieved (1s) or color is retrieved (1c). The retrieval accuracy was estimated by the same classifiers trained on the shape and color map representations (SVM_SS_ and SVM_CC_).

Condition 2 (encode visual + categorical, retrieve visual): shape and color map activations are stored together in the BP along with shape and color labels for each item; either shape is retrieved (2s) or color is retrieved (2c). the retrieval accuracy was estimated as in condition 1.

Condition 3 (encode visual + categorical, retrieve categorical): shape and color map activations are stored in the BP alongside shape and color labels; either shape label is retrieved (3s) or color label is retrieved (3c). The retrieval accuracy of labels was computed by comparing the pre-encoding one-hot representations estimated by the classifiers for each item when it was first classified with the labels reconstructed from the BP.

Condition 4 (encode 50% visual + categorical, retrieve categorical): This was similar to condition 3 except that the encoding parameters for the visual attributes was set at 0.5, to prioritize categorical information over visual. When both shape and color maps are stored as visual information in the BP along with the one-hot coded labels, the visual information was not greatly perturbed (see condition 1 vs. 2 in Table 3 and Figure 12). This is because the one-hot labels are akin to a digital form of information that causes little interference with the stored visual details. It is also evident that retrieving labels result in higher accuracy compared to the visual information when all visual and categorical information are stored in one memory trace specifically for larger set sizes (condition 2 vs. condition 3 in Table 3 and Figure 12). By reducing the amount of visual information stored in memory down to 50%, the effect of set size on retrieving labels becomes smaller (condition 4 in Table 3, and Figure 12). On the other hand, accuracy of storing and retrieving only visual information is sensitive to the number of items (condition 1 in Table 3 and Figure 12). These simulation results match the common finding that people are able to remember several distinct familiar objects that have well-learned categorical labels (i.e., digits or familiar colors) with high accuracy up through approximately 3-5 items, while working memory for specific shape details is more limited (Alvarez & Cavanagh 2004).

**Figure 12.**
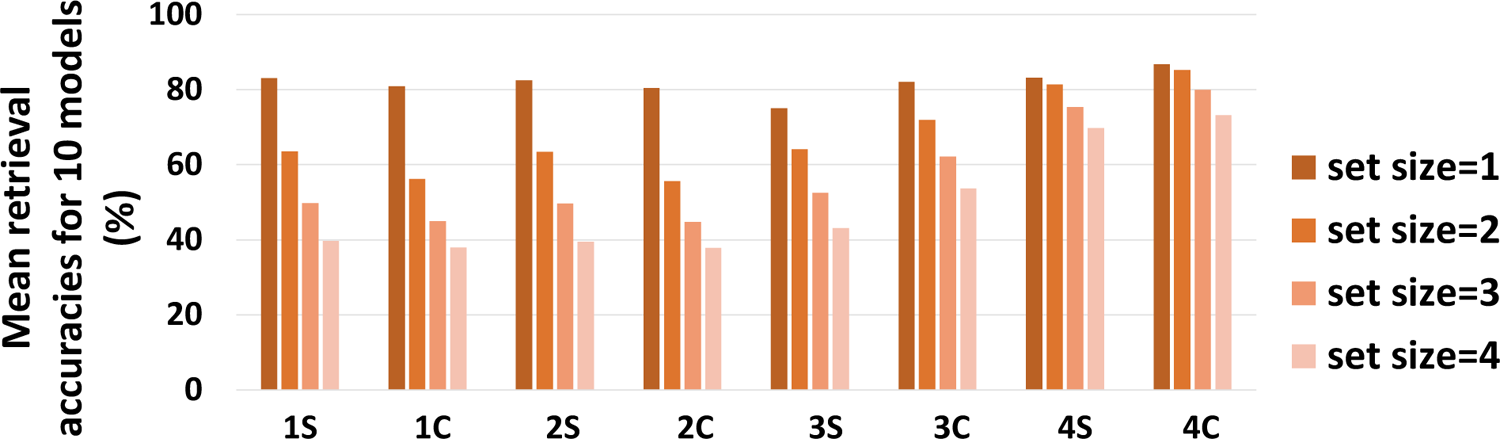
Visualization of Table 3 for mean accuracies of retrieved shapes and color maps (drawn from the classifiers) and labels.

1. 7. *BP binding and content addressability:* Through token individuation, the BP is able to store the attributes of a given stimulus in a combined representation that allows it to link a particular shape to its color. Importantly this allows content addressability (Gorgoraptis et al., 2011), such that if two colored digits are stored in memory using the shape or color representations, memory can be probed by showing just the shape of one of the items and retrieving the token associated with that item. That token can then be used to retrieve the complete representation of the stimulus, including its color. Figure 13 illustrates an example of such binding and subsequent retrieval by a shape cue, and vice versa.

**Figure 13.**
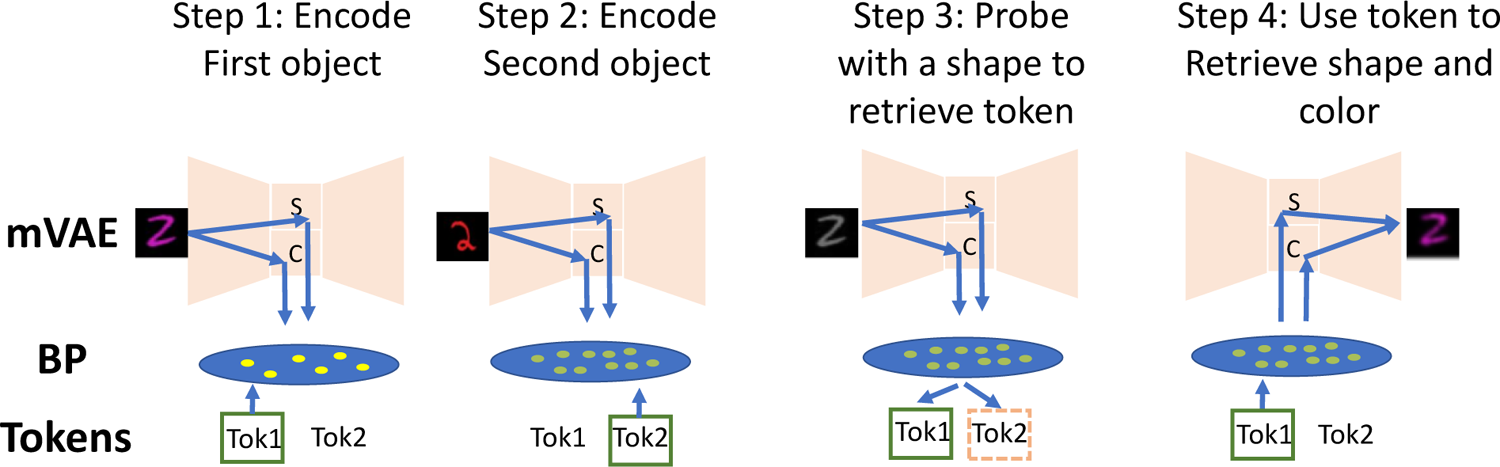
A diagram showing the flow of information during binding. The MLR stores two colored MNIST digits sequentially (step 1and step 2) in the BP. A grey scale shape cue is used to probe and retrieve the corresponding token (step 3). The resulting token is used to retrieve the shape and color of the cued input (step 4). The MNIST digits shown in this figure are not the result of direct simulation, but are just examples to show how binding process occurs.

To test accuracy of binding retrieval, 500 digit-pairs were stored in the BP using the color and shape maps and two tokens. Afterwards, a grayscale MNIST was used as a retrieval cue to determine how often the model successfully retrieved the correct token based on this cue (Figure 13). When the two digits were from two different digit categories (e.g., a “2” and a “3”) the mean classification accuracy across the 10 trained models was 87% (*SE*=.45) with a baseline of 50% chance. We are also able to simulate a case where the model stores two of the same MNIST digit (e.g., two 2’s with a slightly different shape), and then measure if the model is able to correctly retrieve the token associated with one of them. In this case the mean classification accuracy across the 10 trained models is 70% (*SE*= .50), notably worse than when the digits were different but still far better than chance. This is a demonstration of retrieving a cue based on subtle variation in shape between categorically identical stimuli. This capacity is a novel prediction of the model, which is that human WM is able to bind features to an item even when shape differences are subtle and the item category is the same. (see Experiment 4 below for the human data).

1. 8. *More efficient storage of familiar information:* It has been shown that people have higher memory capacity for familiar items drawn from long-term knowledge than novel stimuli (Chen & Cowan,2005; Ngiam, et.al., 2019; Zimmer & Fischer, 2020). The MLR model can show how familiar items are stored more efficiently than unfamiliar ones, and therefore have less degradation of representations in WM as the set size increases. As shown earlier, the BP better encodes the compressed shape and color representations for familiar items (see Figure 9), whereas novel shapes must be encoded from L_1_ and then pass through the skip connection for precise retrieval (see Figure 10). To quantify the memory performance, instead of classifiers we compared the pixelwise cross-correlation of input and retrieved images as the function of set size for familiar and novel^2^ stimuli, such that familiar shapes are encoded from the shape/color maps and novel shapes are encoded from L_1_ and retrieved from the skip connection. The result of the cross-correlations for 500 repetitions are illustrated in Figure 14.

**Figure 14.**
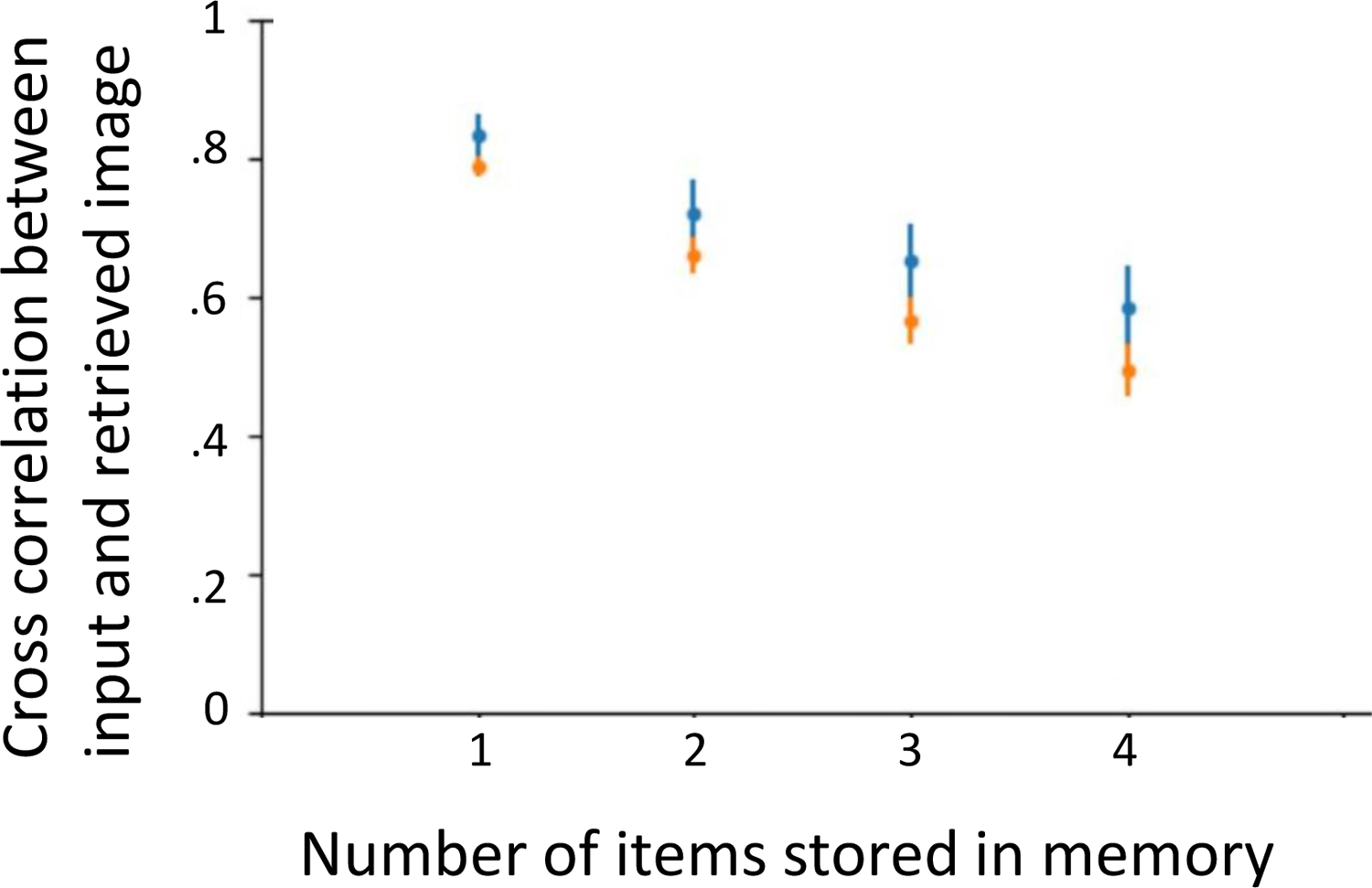
Mean cross-correlation of pixel values for 500 repetitions between input and retrieved images of 10 trained models for familiar (blue) and novel (Bengali, orange dots) shapes across different set sizes Blue dots indicate the correlation for familiar stimuli when the shape/color maps were stored in the binding pool and retrieved via the mVAE feedback pathway. Orange dots indicate the correlation of a novel stimulus when the L_1_ latent was stored and then reconstructed with the skip connection. Note that the reconstruction quality is lower for novel shapes and also that novel reconstructions deteriorate more rapidly as set size increases. The bars stand for standard errors.

As it is shown, the correlation value declines as the set size increases, but more steeply for novel than familiar stimuli. Using cross-correlation, we also measured the memory performance for when familiar items are encoded from L_1_ and retrieved via the skip connection, versus when novel items are encoded from the shape/color maps. The values have been summarized in Table 4. The shape/color map memory retrievals of the novel shapes are ∼15% for all the set sizes, indicating that novel configurations cannot be represented by the highly compressed maps at the center of the mVAE (see also Figure 10 for illustration of memory retrievals from the shape and color maps of Bengali characters). The results also revealed that the L_1_ encoding of familiar shapes and retrieving it via the skip connection yielded a lower performance across all the set sizes compared to encoding of shape and color map representations. Hence, the compressed shape and color representations achieved by training allows for more precise memory representation for familiar shapes, whereas this efficient representation does not exist for novel configurations. Therefore, the model relies on the early-level representations of L_1_ to store novel shapes, which in turn comes at a cost of less precise memory retrievals.

**Table 4.** The correlation values between input and retrieval stimuli as a function of set size

|  | Stimuli type |  |  |  |
| --- | --- | --- | --- | --- |
|  | Familiar |  | Novel |  |
| Retrieval | S/C maps | L1-skip | S/C maps | L1-skip |
| Set size |  |  |  |  |
| 1 | .84 (.03) | .79 (.03) | .15 (.06) | .79 (.01) |
| 2 | .72 (.05) | .69 (.04) | .15 (.05) | .66 (.03) |
| 3 | .65 (.05) | .61 (.04) | .14 (.06) | .56 (.03) |
| 4 | .59 (.06) | .51 (.05) | .14 (.06) | .50 (.04) |

Note. The mean cross-correlation between stimuli and their retrievals for different set sizes across 10 trained models. S/C maps stands for shape and color maps. The values in parentheses are standard errors. The correlation values were measured in cases where the BP encoded the shape and color activations of the novel/familiar stimuli and then the stimuli were retrieved via the feedback pathway (retrieval condition: S/C maps). The correlation values were also measured in cases where the BP encoded the L<sub>1</sub> activations of the novel/familiar stimuli, and then the stimuli were retrieved via the skip connection (retrieval condition: L<sub>1</sub>-skip).

It should be noted that the efficient representation of the shape and color maps is limited to when the model *encodes* the information into memory. In other words, if the images were to be reconstructed from the mVAE without being stored in memory, the L_1_ representations contain slightly more visual detail than the shape and color maps. This is illustrated in Figure 15 for familiar items, in which we computed the correlation values of a single input and its reconstruction when images were reconstructed from L_1_ via the skip connection versus when they were reconstructed from the shape and color maps in the no memory storage condition (left panel). This was compared to memory retrievals from L_1_ and shape/color maps, for which the retrievals from the latter were shown to be more precise (right panel).

**Fgure 15.**
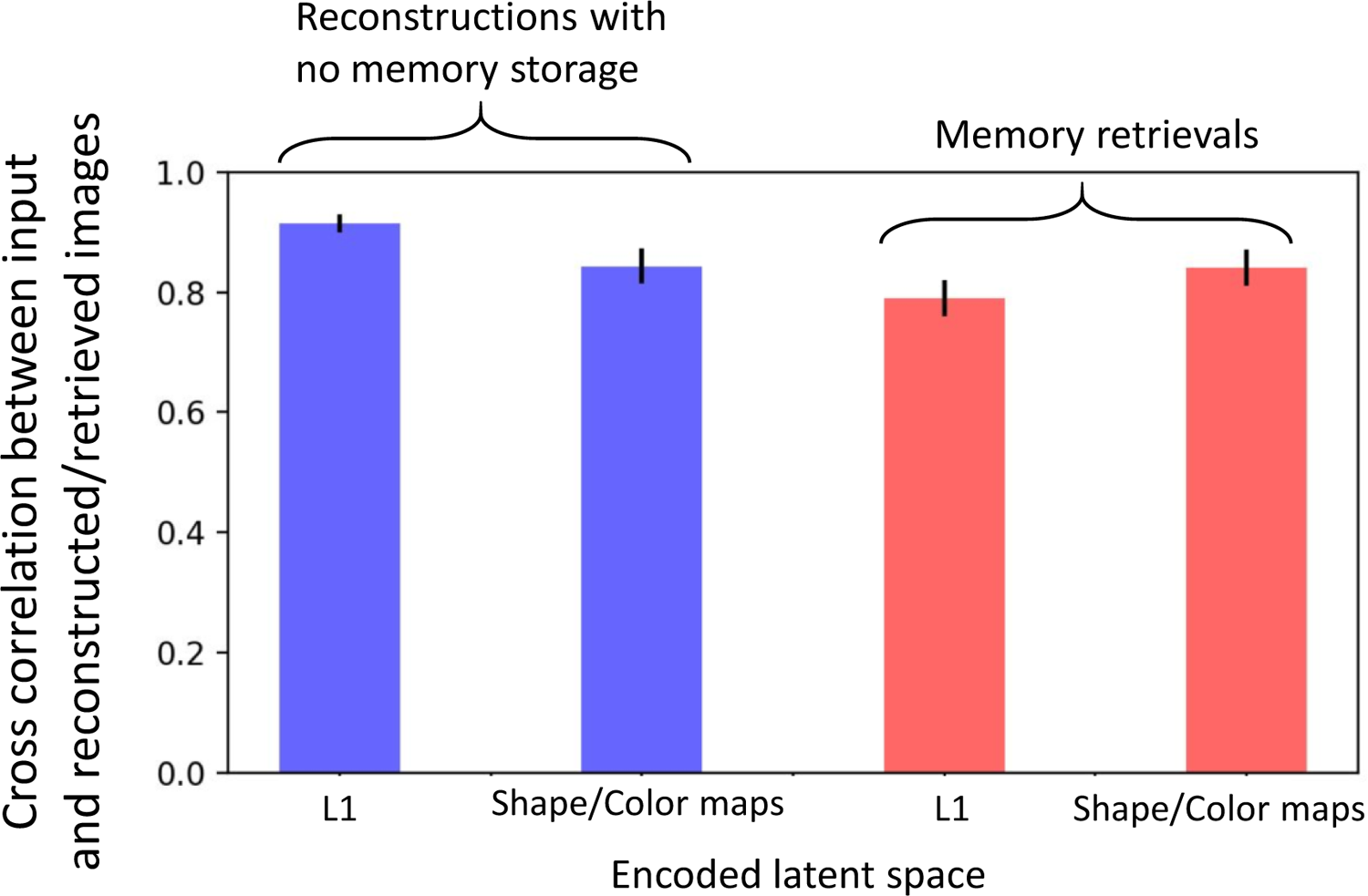
The mean cross correlation values between a familiar input and a reconstructed image for 10 trained model. The blue bars are the mVAE reconstructions of L_1_ via skip connection (left-blue) and the shape/color maps without being stored in memory (right-blue). The red bars are memory retrieval of one item when the L1 representation is stored in the BP and retrieved via the skip connection(left-red) vs. when the shape/color maps are stored in the BP and then retrieved via the decoder (right-red). The error bars represent standard error.

### Empirical Validation of the MLR Model

In partial validation of the model, we provide a series of predictions with empirical tests about the capabilities of working memory in storing visual information. These capabilities were derived from the general properties of the MLR model.

*Prediction 1*: Working memory experiments are typically performed with repetitive experience using the same stimuli that enable participants to develop fine-tuned expectations about the task demand. We posit, however, that WM is typically used without such expectations in daily life and can store mental representations that are useful despite having no expectation of the memoranda or response. The specific prediction is that people can remember the fine-grained shape details of stimuli that they are not very familiar with even in the absence of specific expectations or experience in the task. MLR achieves this result by encoding the L_1_ representations of the Bengali characters, and retrieving them via the skip connection. Cross-correlation of pixel values between the input and the retrieved image was .79 (*SE* = .01) for one item across 10 independently trained models (See Figure 10 for visualizations of memory reconstructions)

*Experiment 1:* 20 Penn State University undergraduates (Mean age = 19.55, 90% female, 20% left-handed) participated for course credit. All the experimental designs were approved by IRB at the Pennsylvania State University, and the scripts are available at this link (https://osf.io/tpzqk/). Participants were shown one Bengali character and were then asked to click on the exact character they remembered seeing from a search array of four Bengali characters (Figure 16). Critically participants were only instructed to pay attention, and were uninformed as to the nature of the ensuing memory question until after viewing the image.^3^ Five Bengali character categories were taken from the stimulus set downloaded from www.omniglot.com, which includes multiple different exemplar drawings of a Bengali character in grayscale. The experiment was developed in Psychopy (v2020.2.2, Peirce et al., 2019) before being translated to JavaScript using the PsychoJS package (v 2020.2) and run online via Pavlovia (Peirce et al., 2019). Each character was presented in the center of a grey screen at size (0.15×0.15 Psychopy height units, a normalized unit designed to fill a certain portion of the screen based on a predefined window size) for 1000ms, followed by a 1500ms delay. After viewing the image, participants were then instructed to click on the image they just saw. Four images including 3 distractors and the target were presented to the participants. On the first trial, non-target answer options were selected from different Bengali characters, and on trial 2 non-target answer options were different exemplars of the same character. Participants were not aware a 2^nd^ trial would occur until after the completed the first. Accuracy scores were considered significantly above chance if a 95% bootstrapped CI (95% bCI) did not include the chance baseline (25%).

**Figure 16.**
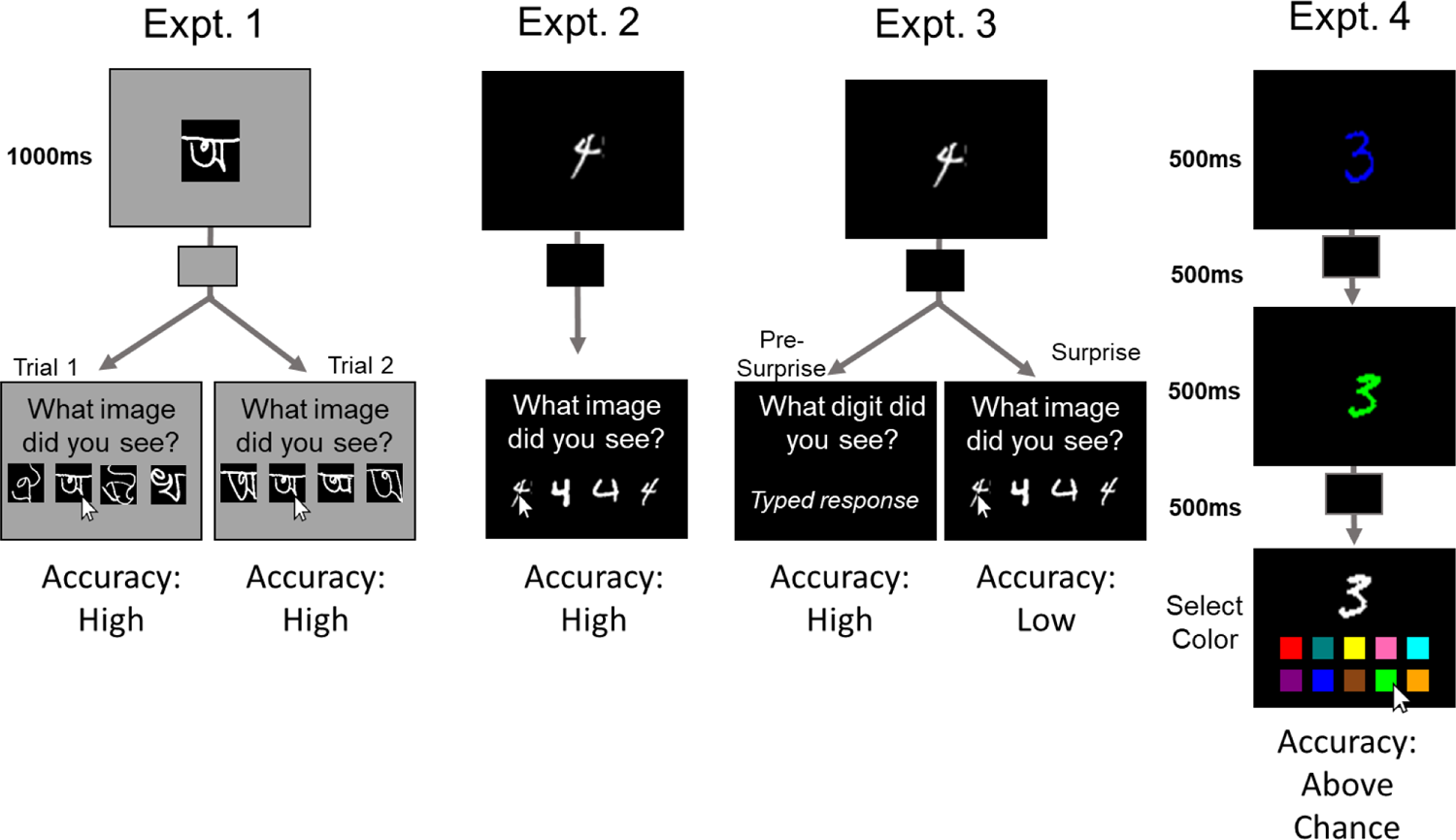
Trial layout for all experiments conducted on human participants. In Experiment 1, participants saw a grayscale Bengali stimulus before being asked to click which image they remembered seeing. The foils presented in the 4-afc varied between trial 1 and trial 2. They were not informed ahead of time that there would be a memory task. Experiment 2 was identical to Experiment 1, except the stimuli used were MNIST digits. In Experiment 3, participants viewed grayscale MNIST and were instructed to type in the category of the image (e.g., type ‘4’ in displayed trial) for 31 consecutive trials before being surprised with a question asking them to click on the exact MNIST exemplar they remembered seeing. In Experiment 4, participants were instructed to remember the color-exemplar pairing of MNIST digits, before being cued with the specific exemplar and asked to click on the color that exemplar was. The key behavioral result is summarized below each condition, see text for details.

*Experiment 1 results:* Though participants were shown novel targets and given no instruction to remember the Bengali character presented to them, the mean accuracy on the very first trial was 95%, 95% bCI [85%,100%]. Participants were also highly accurate on the second trial which asked a more difficult question by requiring them to find the target image from the same category, *M* = 90%, 95% bCI [75%,100%].

*Prediction 2*: When people create a memory for a familiar stimulus, they can also have a memory for the specific shape of that stimulus, as well as being able to categorize it. This is particularly true when there is no specific expectation as to what response will be necessary. MLR predicts this by storing multiple codes of a given visual stimulus in one memory trace.

*Experiment 2:* A new sample of 20 Pennsylvania State University undergraduates (Mean Age = 18.6, 90% female, 5% left-handed) participated in this online experiment for course credit.

Participants viewed one grayscale MNIST digit image (3, 4, 6, 7, and 9) on a black background before being asked to click on the exact image they remembered seeing (Figure 16). Again, participants were not informed there would be a memory task^4^. Thus, the first trial served as an unexpected memory test. Non-target options were exemplars from the same digit category (e.g., they saw four different instances of the digit 3, one of which was an exact match to what they had just seen; see Figure 16). Participants completed 5 trials in total, with a new digit category shown on each trial (i.e., digit categories were never repeated within an individual). All other components of Experiment 2 were identical to Experiment 1.

MLR simulates the lack of expectation via setting the encoding parameters of the shape and color maps to 1.0. and encoding both features as well as the categorical labels into memory. We replicated simulation 6 (Table 3, condition 2s) except with grey scale stimuli. The mean decoding accuracy of the retrieved shape across 10 trained models was 83% (*SE* = .43).

Results: Overall, the mean accuracy of reporting the MNIST exemplar without being specifically instructed was 85% (95% bCI [60%, 95%]) on the very first trial. Accuracy on subsequent trials was 85% (95% bCI [70%,100%]), 90% (95% bCI [75%, 100%]), 100%, and 100%. Thus, people could remember the shape of a familiar stimulus even when there was no expectation to report on the its shape and improved to perfect accuracy with a small amount of experience. This supports our assumption that even highly familiar stimuli are encoded at the specific shape level in the absence of expectation of what specific question will be asked.

Prediction 3: Our next prediction is that expectation can tune representations for highly familiar objects to represent categorical information at the expense of visual information. This will be tested by asking subjects to repeatedly report the identity of a digit, ignoring its specific shape, and then giving them an unexpected question about its shape after 30 trials. Thus, the same question about visual shape that could be easily answered in Experiment 2 should be hard to answer after memory encoding settings have been tuned to exclude visual details.

This prediction stems from the fact that MLR has modifiable parameters controlling the relative contribution of shape vs one-hot categorical representations as memories are constructed. We modified the parameters similar to simulation 6 (Table 3, condition 4) except that only 20% of shape information is stored in memory alongside the categorical label. The classification accuracy of the decoded shape information demonstrated 24% (SE=1.6) shape accuracy, whereas the accuracy of retrieving the label was 84% (SE =.6). It should be noted that the ceiling accuracy of labels is constrained by the classifier accuracy, which is about 85% and thus lower than human performance in identifying an MNIST digit, which is near 100%.

Experiment 3: A new sample of 20 Pennsylvania State University undergraduates (Mean Age 18.8, 95% female, 5% left-handed) participated in this online experiment for course credit. The paradigm (Figure 16) resembles that used in standard Attribute Amnesia studies (Chen & Wyble, 2015). Participants viewed a grayscale MNIST digit (from any digit category 0 through 9), and were instructed to report the category of the image by typing the respective digit on the keyboard^5^. This task was repeated for 50 trials before participants were asked a surprise question on Trial 51: instead of identifying the image category, they had to select the specific category exemplar they remembered seeing (i.e., which specific “2” among a 4-AFC array of MNIST “2s”). On the surprise trial, participants reported the specific shape of the digit they just saw by clicking on the image that matches the target^6^. The display response consisted of the target and 3 distractors from the target category but with different shapes (see Figure 16).

Participants then completed 9 more exemplar identification trials (termed *control* trials). Significance for accuracy changes on the surprise trial was assessed by comparing surprise trial accuracy to accuracy on the 1^st^ control trial via a permutation test (10,000 iterations). All other parameters of this study were identical to Experiment 2.

Results: The mean accuracy of identifying the target was 97% during the 50 pre-surprise trials. However, on the surprise trial, the accuracy of identifying the exact shape of the presented stimulus was 15%, 95%bCI [0,30%]. On the very next trial, when participants had an expectation to report such information, the accuracy of reporting the shape of the digit elevated to 100% on the very next trial. The difference between performance on the surprise and first control trials was significant as determined via a permutation test, p <.0001. This demonstrates that memory representations can be tuned to represent largely categorical information, with minimal specific shape information.

Prediction 4: Perhaps the most striking capability of WM that is unique to MLR among competing models is the ability to bind two specific colors to two different shapes, even when the two shapes are from the same category (binding accuracy: *M*=69.92%, *SE*= .50). This follows from the fact that tokens link shape map information to color map information.

Therefore, when shown two different instances of an MNIST digit of a given category with different colors, participants should be able to report which color was bound to which specific shape (See Figure 13)

Experiment 4: A sample of 20 participants (Mean Age 21.9, 45% female, 15% left-handed) were recruited from the online website Prolific. On each trial (Figure 16), 2 MNIST exemplars from the same digit category were presented sequentially to the participant. Each exemplar was randomly colored from a list of 10 options (Red, Green, Blue, Pink, Yellow, Orange, Purple, Teal, Cyan, and Brown), and colors did not repeat within a trial. Each digit was visible on screen for 500 ms, with a 500 ms ISI between exemplars and a 500 ms delay between the second exemplar and the response screen. One of the exemplars (counterbalanced across trials) was then presented to the participant in grayscale, and participants were instructed to click on the color that was paired with this exemplar (10-afc; chance = 10%). Unlike in previous experiments where no instruction was given, participants *were* explicitly instructed to remember the color-shape pairing^7^OBJ.

Results: Participants were able to remember which color was linked to which specific MNIST exemplar. Overall, Participants correctly reported the target color 81.5% of the time^8^, 95% bCI [75.5%, 87.25%], with swap errors (reporting the color of the other MNIST digit) occurring on average 9% of the time, 95% bCI [4.75%, 14%]. Importantly, participants were capable of completing this task on the first trial, as 17 of 20 participants (85%) reported the correct color on trial 1, 95% bCI [70%, 100%]. This shows that the ability to utilize one attribute of a bound representation to retrieve another attribute is a general capability of WM, rather than a specific capacity that emerges through training.

### General Discussion

The MLR model provides a plausible account of rapid memory formation that utilizes a limited neural resource to represent visual and categorical information in an active state. The model mechanistically explains how WM representations could build on long-term knowledge traces, capitalizing on visual experience to store familiar items more efficiently. Using a generative model such as a VAE, we were able to build a knowledge system based on synaptic plasticity. The VAE layers also share similarities with the visual ventral stream and can represent complex shapes with compressed representations. We showed that when images are drawn from MNIST or f-MNIST datasets as familiar stimuli, the visual knowledge provided the WM with the item’s compressed representation at its later levels or its categorical label. Whereas, for novel shapes the model could leverage generic representations at early layers to encode them into WM. Subsequently, we demonstrated that the advantage of storing compressed format of known shapes is having less interference between items compared to when early level representations of novel shapes^9^ are stored in memory(Figure 14). Moreover, we showed attribute binding for individual items by encoding 2 digits with different colors in WM, and cuing one of the shapes to retrieve the whole item. Finally, we provided empirical evidence of the basic principles of the model, showing that naïve subjects can retrieve the specific shape of both novel and familiar stimuli and can bind attributes to presented shapes.

#### Addressing the functional capabilities of Working memory

There is a tremendous amount of literature on the functional capabilities of working memory. Thus, it is impossible to address all of it here. As detailed below, there are specific and important aspects of the memory literature that we are unable to address due to the scope of the model, but the model is able to address high-level requirements of a working memory model as provided by Oberauer (2009). Here we describe them with reference to MLR.

1. *Build new structural representations*: This refers to the ability to quickly link or dissolve representations that bind parts of existing representations together into novel configurations. This is demonstrated by the ability of MLR to form representations of novel spatial arrangements of line segments (i.e., Bengali characters) extracted from the L_1_ latent.
2. *Manipulating structural representations*: This refers to the ability to access information that is currently stored in memory and to implement cognitive operations on it. MLR does not represent cognitive operations, but it has tunable parameters that control the flow of information to determine what specific attribute(s) or labels should be encoded into WM.
3. *Flexible reconfiguration*: This refers to findings that WM is a general-purpose mechanism that can be reconfigured. This flexibility is at the heart of MLR’s mechanism for adjusting which latent spaces are projected into the binding pool according to task requirements. For instance, novel configurations are stored using a different latent than familiar shapes. Moreover, changing the encoding parameters does not require adjusting gradients, but is instead a modulation of transmission along a given pathway.
4. *Partially decoupled from long-term memory*: WM must be able to store and retrieve information in a way that is distinct from information stored in long term memory. The binding pool exhibits exactly this property by creating active representations that are separate from the latent spaces embedded in the visual knowledge.
5. *Draws on long-term memory:* The primary function of MLR is its ability to build efficient memories on existing long-term knowledge representations. The BP in MLR can use compact latent spaces for stimuli that the model had training experience with.
6. *Transferring useful information into long-term memory*: It must be possible to convert or “train” WM representations into long-term memory representations. This capability is enabled by the generative aspect of MLR. Memories for novel stimuli can be reconstructed at the earliest levels of input and can then be used to drive learning mechanisms at any subsequent latent representation. In other words, memory consolidation could occur by regenerating remembered representations and then using those to drive perceptual learning (or gradient descent in an artificial neural network).

Another requirement emphasized by Norris (2017) is the need for a short-term memory system to explain how novel visual stimuli can be remembered, which MLR can do by virtue of reconstructing the L_1_ latent with the aid of BP. Norris also discussed extensively the distinction between copies of information and pointers. In MLR, memory encoding is a combination of the two. When a latent space is projected into the BP, it is not a literal copy of the original pixelwise stimulus. Instead, the BP stores a copy of the representation in a latent space, and thus requires that latent space, and the supporting circuitry in the feedback pathway to reconstruct the information at the pixel level.

More detailed constraints on WM can be found in a comprehensive list of empirical benchmark effects (Oberauer et al. 2018). Future iterations of computational models such as MLR with a larger scope and a more direct simulation of time, could take advantage of such data to constrain more detailed models.

#### Comparison of approaches

As mentioned in the introduction, there are several major frameworks for thinking about the storage of information in WM in relation to long-term knowledge. Separate-store accounts imply a delineation between perceptual systems, long-term memory, and the WM system (Atkinson & Shiffrin 1968; Baddeley & Hitch, 1974; Baddeley, 2000). This separation is consistent with the notion of prefrontal cortex as a substrate for WM as discussed in the electrophysiological literature (Goldman-Rakic,1995; Miller, Erickson & Desimone, 1996). These accounts can be contrasted with embedded process accounts (Cowan 1988, 1999, Cowan, Morrey & Benjamin-Naveh 2018; Oberauer 2009, Teng & Kravitz, 2019) in which there is less of a distinction between WM and other cognitive systems such as long-term memory.

The MLR has a binding pool for storage that is distinct from the perceptual system and visual knowledge and therefore is more in line with the classical separate-store account. However, the MLR model could in principle be modified to resemble an embedded process account by placing small binding pools within each of the processing layers. This example illustrates the value of implemented models, as they provide a computational intuition for contrasting accounts, even in the absence of explicit implementation. Making such a modification to MLR, without losing its ability to bind features together would require that the tokens link to each of the small binding pools. Moreover, such an account would not exhibit very much interference between different levels of representation given that they were represented in different cortical areas (i.e., one could store familiar and unfamiliar objects together with minimal interference).^10^

Another distinction in the space of WM models is whether there are unitary or separate memory storages for different types of information (i.e., visual and non-visual). MLR makes no distinction between the storage of information in that it encodes categorical and visual information in one memory trace. While auditory representations are not included in MLR, it would be possible to project latent representations of phonological information into the binding pool as well. However, the model could be amended to incorporate a distinction between visual and non-visual information by adding a second binding pool. We are persuaded by Morrey (2018) that the evidence for such a distinction is not clear cut and thus are proposing the simpler and parsimonious unitary account. Developing computational theories such as this is a means to advance these debates by providing more tangible implementations of these theoretical positions.

#### Advantages of the binding pool architecture in MLR

Given these possible implementations, we propose that a key advantage of clustering neural activity associated with memory into a binding pool of general-purpose storage neurons is that higher-order processes have a straightforward path to control those representations, allowing them to be sustained, deleted, or instantiated into constituent cortical areas with a relatively small amount of circuitry. In the context of MLR, it is harder to imagine how a centralized executive control system could exert control over an embedded memory system, since those control circuits would have to synchronously infiltrate a large number of cortical areas in order to reactivate, manipulate or extinguish those memory representations. Binding information between different features within distinct objects is also simpler to implement in a binding pool architecture as demonstrated here because the information is physically clustered in a well-defined population of neurons. Thus, the arguments in favor of an account like MLR is that it places less demands on the interplay between functional networks. On the other hand, it could be that synchrony mechanisms could play a role in aligning tokens with an anatomically distributed binding pool in the absence of highly specific anatomical connections (Miller, Lundqvist & Bastos, 2018).

Finally, perhaps the strongest argument in favor of having a BP mechanism is that BP makes storing episodic representations into long-term memory fairly straightforward. By building memories of the binding-pool activity state, the hippocampus can store a copy of the most relevant information at the time the WM was constructed. Thus, a snapshot of the binding pool state provides a combined memory trace of the information that was deemed important or task relevant at the time that trace was constructed.

#### Candidate regions for the binding pool

The binding pool is instantiated here in a mathematically idealized form of a single pool of neurons that is linked with an undifferentiated connection to a wide range of cortical areas. This is the most naïve solution with no training or optimization of the weights between the latents and the BP, and it is of course more likely that a biological instantiation would have differentiation to represent information more efficiently. Thus, the simple random connectivity we use here should not be taken as a strong assumption that there is no learning of this weight matrix in the real system.

As far as physical realizations the claustrum is an example of a brain area that has the potential cortical connectivity to support a domain general binding mechanism (Crick & Koch 2005; Fernández-Miranda, Rhoton, Kakizawa, Choi & Álvarez-Linera, 2008). It is also tightly interconnected with the medial entorhinal cortex (in mice, Kitanishi & Matsuo 2017) which would allow binding pool representations to be stored in the hippocampus as episodic traces. It also allows remembered binding pool representations to be reconstructed from the hippocampus and then projected down to constituent areas. Rhinal areas leading into the hippocampus could also serve this role, thus enabling transfer of information from the binding pool into more durable episodic memory representations. However, rhinal areas are generally not directly connected to lower order sensory areas, which would make it more difficult to store representations from very early areas.

Another possibility is that the functionality of the binding pool could be distributed through parts of the basal ganglia to provide a highly flexible form of working memory that incorporates top-down control circuitry to determine what information is stored at any point (O’Reilly & Frank 2006). The involvement of prefrontal areas in WM has long been suggested by single unit work in monkeys (Goldman-Rakic,1995). In particular, the mixed selectivity of neurons in the prefrontal cortex across tasks (Rigotti, et al 2013) and sequential order (Warden & Miller 2007) is emblematic of how neurons in the binding pool would exhibit different tuning properties depending on a particular configuration of cognitive processing strategies (see Wyble, Bowman & Nieuwenstein, 2009 for discussion on this). Thus, a system in which the prefrontal cortex embodies the token control architecture, while the binding pool resides in areas like the basal ganglia or the claustrum are potential candidates for a binding pool.

#### Limitations

MLR is not intended as a complete model of working memory as there are many functional, empirical and computational aspects that have not yet been considered. Their omission is not intended to signal that they are unimportant, but rather is an admission that a formal implementation of a cognitive function so flexible as WM is beyond the scope of any single paper or even career (See Cowan 2001, Barrouillet et al.,2009, Logie Camos & Cowan 2021, Oberauer, 2009; Oberauer et al. 2018; Schneegans &Bays, 2019; for extensive discussion on other aspects of WM) Rather, the MLR is intended as a nucleus of a storage mechanism to store memories in a way that is linked to visual knowledge and that is extensible to a broader range of empirical phenomena and capacities. Here are several aspects of the model that have not been considered but are likely crucial for a more comprehensive account of WM.

*Space*: MLR in its current implementation has no ability to represent different spatial locations across the visual field, as the input space is only 28×28 pixels and one stimulus fills most of it. Adding biologically realistic spatial location as a dimension to an autoencoder such as the mVAE is feasible in principle, although it would require additional layers and complexity in terms of connectivity gradients across foveal and parafoveal areas. We consider such improvements to be an important next step in the development of models such as this. The binding pool as a storage mechanism is generic with respect to information content and thus should be adaptable to any model containing latent spaces.

*Time:* For simplicity, there is no explicit representation of time in MLR, which makes many empirical phenomena inaccessible. For example, primacy and recency gradients, the time course of memory consolidation and decay of memory over time are not within the scope of the model. Future work could address this with a more detailed neural model.

*Attention and central executive:* For simplicity, we assume that control circuits determine when information is encoded, or reconstructed, and also tune the encoding parameters so that the most relevant latents for a given memory trace are stored. Attention is thereby implemented in parameters that are tuned to produce efficient encoding, but no attention is required for maintenance of information in the current implementation. Attention would also be required for managing the flow of information between different areas of the model (e.g., encoding from latents into the BP, retrieving information from the BP, or erasing WM; see appendix for further discussion of this point).

## Conclusion

Working memory has been at the center of cognitive psychology and neuroscience research for decades. It is evident that this mental construct is inseparable from our previously learned knowledge, as it has been stated in many of the existing theories (Baddeley, 1992; Cowan, 1999). However, without an actual implementation of a WM model that is coupled with visual knowledge, contrasting and evaluating the existing theories seems to be very challenging, if not impossible. The MLR model, constrained by computational, biological and behavioral findings is the first attempt to shed light on the computational mechanisms of WM in relation to visual knowledge, while highlighting the aspect of familiarity in storing and retrieving visual information. We believe implementing models as such are valuable resources for generating intuitions about the rapid formation of representations in working memory.

## Acknowledgements

We would like to thank John Collins, Joyce Tam, Chloe Callahan-Flintoft and Pooyan Doozandeh for their helpful comments during the preparation of this manuscript. This work was supported by NSF grant 1734220 and Binational Science Foundation grant 2015299.

## appendix

### Colorized MNIST and Fashion MNIST

The original datasets of MNIST and f-MNIST are in grey color. Each image in the dataset was randomly colorized by 10 prototype colors – red, blue, green, purple, yellow, cyan, orange, brown, pink, teal – with color values being [[0.9, 0.1, 0.1], [0.1, 0.9, 0.1], [0.2, 0.2, 0.9], [0.8, 0.2, 0.8], [0.9, 0.9, 0.2], [0.1, 0.9, 0.9], [0.9, 0.5, 0.2], [0.6, 0.4, 0.2], [0.9, 0.7, 0.7], [0.1, 0.5, 0.5]].

The color of each image was chosen by first selecting a prototype color and then adding random variation to each of the RGB channels from the range [-.1, .1].

### Modified VAE

Our innovation in modifying the VAE was using 3 objective functions to train the shape map, color map and the skip connection. All the objectives were derived from Equation 1s that is used for VAE. The explanation for variables and parameters of the following Equation can be found in Kingma & Welling (2013).

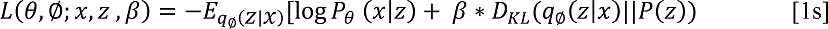

*Skip objective function:* minimizes the Equation 2s, where the first term represents cross entropy and the second term represents the KL divergence for input x and the bottleneck variable of z_s_ for shape and z_c_ for color map. z= z_s_ + z_c_ . The mathematical explanation of all the variables can be found in Kingma & Welling, 2013. We adopted β = .25.

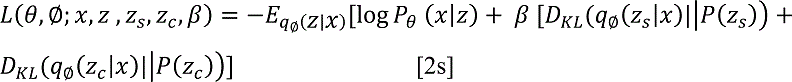

Shape objective function: This function converted images into grey scale images by averaging across the three RGB channels. Then the following objective was minimized with β = .5.

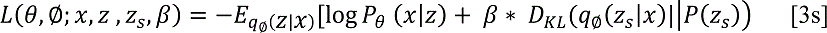

While the shape map was being trained, color map and skip connection were detached.

Color objective function: This function computes the maximum color value of rgb channels for each image and converts the image to those value. That results in replacing each image with color patches of highest value. Then, it minimized the Equation 4s with β = .5.

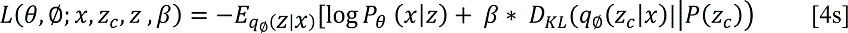

While the color map was being trained, shape map and skip connection were detached. *BP Connectivity*

The binding pool weight matrix was generated via randomly normally distributed values (mean 0, std dev= 1.0) and are randomly re-generated for each simulated trial. However, they remain fixed each time that the binding pool function is called.

### Token Connectivity

Each token is connected to a random set of 40% (i.e., 1000) of total nodes (i.e., 2500). This means that when a given token is active, that subset of BP nodes it is connected to can be used to store and retrieve information, the remaining BP nodes will still hold their activation state, but can neither be encoded to, or retrieved from. The subset of BP nodes associated with each token overlap with one another so that for any given token, 40% of its nodes overlap with any other token. As a result, with increasing the number of tokens stored in memory, the likelihood of interference between objects increases due to the overlap between token connectivity to the BP. There is no limit on the number of tokens, but the binding pool is assumed to be fixed in size.

### Control signals for encoding and retrieval

As in the simpler binding pool architecture of Swan & Wyble (2014), it is assumed that there are control signals that determine when information is encoded into or retrieved from the binding pool. These would correspond to commands from a central executive and take the form of volumetric neuromodulation or suppressive gating signals that can determine whether afferent or efferent connections are active at any given point. These control signals are implemented with procedural code for simplicity.

### SVM classifiers

SVM classifiers were used to determine the information represented in shape and color maps as well as estimating categorical labels. SVMs were imported from the scikit-learn library as radial basis functions (kernel= ‘rbf’) with the decision function parameters to be C=10 and gamma=’scale’ respectively.

## Footnotes

^1^Tokens could be implemented either by enabling encoding in their own BP nodes through excitation of gate nodes, or alternatively by disabling encoding in the other BP nodes through suppression of gate nodes. These methods are functionally equivalent.

^2^Due to the limited number of images for Bengali characters as novel shapes, we augmented the data by doing slight rotation (10° rotation) and random crop with padding =8 on the 6 characters. This enabled us to do the permutations test for measuring cross-correlation.

^3^The exact instructions for the experiment were provided separately on each trial, to minimize the potential of participant’s predicting what they may need to remember during this experiment. Thus, the instructions occurring before trial 1 were as follows: “Thank you for participating in this experiment. You will be completing two separate experiments! This 1st experiment will be a very short, ONE TRIAL experiment where we show you some visual information. Because there is only one trial we need your full attention, as you only get ONE SHOT. So keep your eyes on the fixation cross before the stimulus appears. Press the SPACEBAR when ready to begin.” The instructions following trial 2 were as follows: “”That concludes our first experiment! We will now begin the 2nd, equally fast ONE TRIAL experiment. We will show you some new visual information. Again, we need your full attention, as you only get one trial. Press the SPACEBAR when ready to begin.”

^4^The specific instruction was: “This experiment will be a very short experiment where we show you some visual information. Because it is short and each of the 5 trials are unique, we need your full attention right from the start. Keep your eyes on the fixation cross before the stimulus appears. Press the SPACEBAR when ready to begin.”

^5^The exact instructions were as follows: “In this task, you will be presented with an image of a digit. Your task is to identify which digit it is. When asked to do so, using the number keys on the top of your keyboard (NOT the number pad), press the key of the number you remember seeing.”

^6^The exact instructions presented on the surprise trial were as follows: “This is a surprise memory test! Try to remember the digit you last saw? What was its specific shape? Click the image that matches the image you just saw”

^7^The exact instructions were as follows: “In this task, you will be presented with 2 images of digits in different colors. The digits will be the same (i.e. two 4s) with different shapes. Please remember which shape is linked to which color. You will then be asked to report the color of 1 of the 2 digits. You will be reshown one of the twp previously seen shapes, and when asked to do so, please click on one of the 10 color choices that was paired with that shape.”

^8^Target presentation order had no influence on accuracy. Participants were as accurate at reporting the target’s color when the first exemplar served as the probe (M = 84%) compared to the second (M = 79%), permutation p-value = 0.49.

^9^It is important to clarify that in reality we cannot have a purely novel stimulus, as any shape can be decomposed into elements that we are more or less familiar with. For instance, an unfamiliar Chinese character can be remembered as lines and strokes or even objects that one has encountered with in her lifetime.

^10^Other embedded process accounts as described by Cowan (2001) are able to explain cross-dimensional interference as an attention limitation and it would be useful to develop implementations of those accounts to compare their functional limitations to models such as MLR.

